# A spatial code for temporal cues is necessary for sensory learning

**DOI:** 10.1101/2022.12.14.520391

**Authors:** Sophie Bagur, Jacques Bourg, Alexandre Kempf, Thibault Tarpin, Khalil Bergaoui, Yin Guo, Sebastian Ceballo, Joanna Schwenkgrub, Antonin Verdier, Jean Luc Puel, Jérôme Bourien, Brice Bathellier

## Abstract

The temporal structure of sensory inputs contains essential information for their interpretation by the brain^1–9^. Sensory systems represent these temporal cues through two codes: the temporal sequences of neuronal activity and the spatial patterns of neuronal firing rate^3,7,10–20^. However, it is still unknown which of these two coexisting codes causally drives sensory decisions^3,10,20,21^. To separate their contributions, we designed an optogenetic stimulation paradigm in the mouse auditory cortex to generate neuronal activity patterns differing exclusively along their temporal or spatial dimensions. Training mice to discriminate these patterns shows that they efficiently learn to discriminate spatial but not temporal patterns, indicating that spatial representations are necessary for sensory learning. In line with this result, we observed, based on large-scale neuronal recordings of the auditory system, that the auditory cortex is the first region in which spatial patterns efficiently represent temporal auditory cues varying over several hundred milliseconds. This feature is shared by the deep layers of neural networks trained to categorise time-varying sounds. Therefore, the emergence of a spatial code for temporal sensory cues is a necessary condition to associate temporally structured stimuli to decisions. We expect this constraint to be crucial for re-engineering perception by cortical stimulation.

## Main text

Many stimuli which drive selective behavioural decisions, such as phonemes and vocalisations^5,10^, tactile textures and shapes^7,20^ or the coherent motion of a moving animal^4^ result from temporally evolving sensory inputs. In the brain, this temporal structure is associated with changes both in when neurons fire and which neurons fire. On the one hand, temporal stimuli drive neuronal activity sequences, observed throughout the visual^3,11^, auditory^12,13,22^, tactile^14,15^ and olfactory^6,16^ systems, including sensory cortex. In single cortical neurons, these activity sequences carry information which is not available in the neuron’s mean firing rate^7,11,13,15,16^. On the other hand, several studies have established that the time-averaged firing rate of many neurons is tuned to specific temporal cues, such as the speed or direction of motion in visual stimuli^9,23^, the dynamics of tactile contacts^14,19^ or amplitude and frequency modulations in sounds^17,18,22,24,25^. This tuning generates a spatial representation of the temporal structure of sensory inputs^10,20^ based on the identity of the set of activated neurons. Although these spatial representations do not necessarily form anatomical maps and can be widely distributed across a sensory area, they constitute a code for temporal sensory cues that depend on the neuronal space rather than on time. The functional importance of these spatial and temporal codes for behaviour is a long standing issue in sensory neuroscience given that they coexist at all levels of sensory systems including the cortex^10,11,15–17,19,20,22,26^. Therefore, direct manipulation of neural activity patterns in space and time is necessary to evaluate which code is actually deciphered by downstream areas to drive perceptual decisions and behavioural output. While activation of spatial patterns in sensory cortex has been shown to drive discriminative behaviour^27–30^, the role of temporal cues has so far only been causally addressed at the level of sensory receptor neurons^26,31^ or of peripheral sensory networks^32^. It is therefore unknown whether, at the cortical stage, temporal codes are exploited by the downstream motor centres within which associations between stimuli and behavioural decisions are learnt^33^.

### Engineering spatial and temporal codes

To address this question, we engineered optogenetically-driven activity patterns in the auditory cortex (AC) that can be distinguished either specifically from their spatial structure or from their temporal structure. We focused on the auditory system because natural sounds contain rich temporal information^2^ that is at the basis of speech recognition^5^ and influence several perceptual properties important for sound identification and characterization such as timber or loudness^1,34^. We used Emx1-Cre x flex-ChR2 mice expressing channelrhodopsin in a large population of pyramidal neurons and optogenetically activated a low (A spot) and a high (B spot) frequency region of primary AC using light spots delivered with a video-projector through a chronic cranial window^27,29^ (**Fig. 1a, Extended Data Fig. 1a-d**). To generate spatially distinct but temporally identical neural patterns, we stimulated either the A or B spot with the same train of ten pulses at 20 Hz (**Fig. 1b**). Generating spatially identical but temporally distinct patterns is more challenging due to two confounding phenomena. First, cortical neurons adapt their firing rate to stimulation frequency^35^ and adaptation varies across neurons. This leads to different firing rates for different neurons, thus generating spurious spatial patterns. We therefore excluded temporal stimulations differing by stimulation frequency and focused on temporal cues based on the relative timing of the stimulation of the two spots^36^ (**Fig. 1b**). However, based on earlier observations^26,31,37^ and neuronal simulations (**Extended Data Fig. 2a-c**), we reasoned that a recurrent circuit, like the cortex, can convert relative timing cues in the stimulation into spatial patterns in the neuronal population (**Fig. 1c**), which would prevent us from generating purely temporal activity patterns in the network. To quantify these effects, we used silicon probes in awake mice to record the light-evoked activity during optogenetic stimulation of the A and B spots, while shifting the delay between each spot stimulation (**Fig. 1d-f**). Stimulation of A or B alone elevated the time-averaged firing rate in different sets of neurons (**Fig. 1e**), hence producing distinct spatial patterns. Optogenetically driven firing rates were in the same range as those naturally evoked by sound presentation (**Extended Data Fig. 2d**). Stimulating the A and B spots successively in the two possible orders (A-B or B-A) elevated time-averaged firing rates similarly, suggesting much weaker spatial information (**Fig. 1e**). To verify this, we used a population decoder measuring if it is possible to discriminate on a trial-by-trial basis between A-B and B-A sequences only based on the time-averaged firing rate of recorded neurons. We found that this spatial decoder could discriminate temporal stimulation order above chance if the interval between A and B stimulations was lower or equal to 13 ms, but could not for an interval of 25 ms (**Fig. 1f**). Hence, consistent with synaptic interactions integrating over the short membrane time constant of cortical neurons *in vivo*^38,39^, only stimulus sequences separated by a sufficiently long time interval generate cortical network patterns that contain robust temporal information (quantified in **Extended Data Fig. 2e**) but lack spatial information (**Fig. 1f**). We therefore selected the successive activations of A and B separated by 25ms and in opposite orders as a protocol to assess discriminability of purely temporal patterns in a behavioural task.

**Fig. 1.**
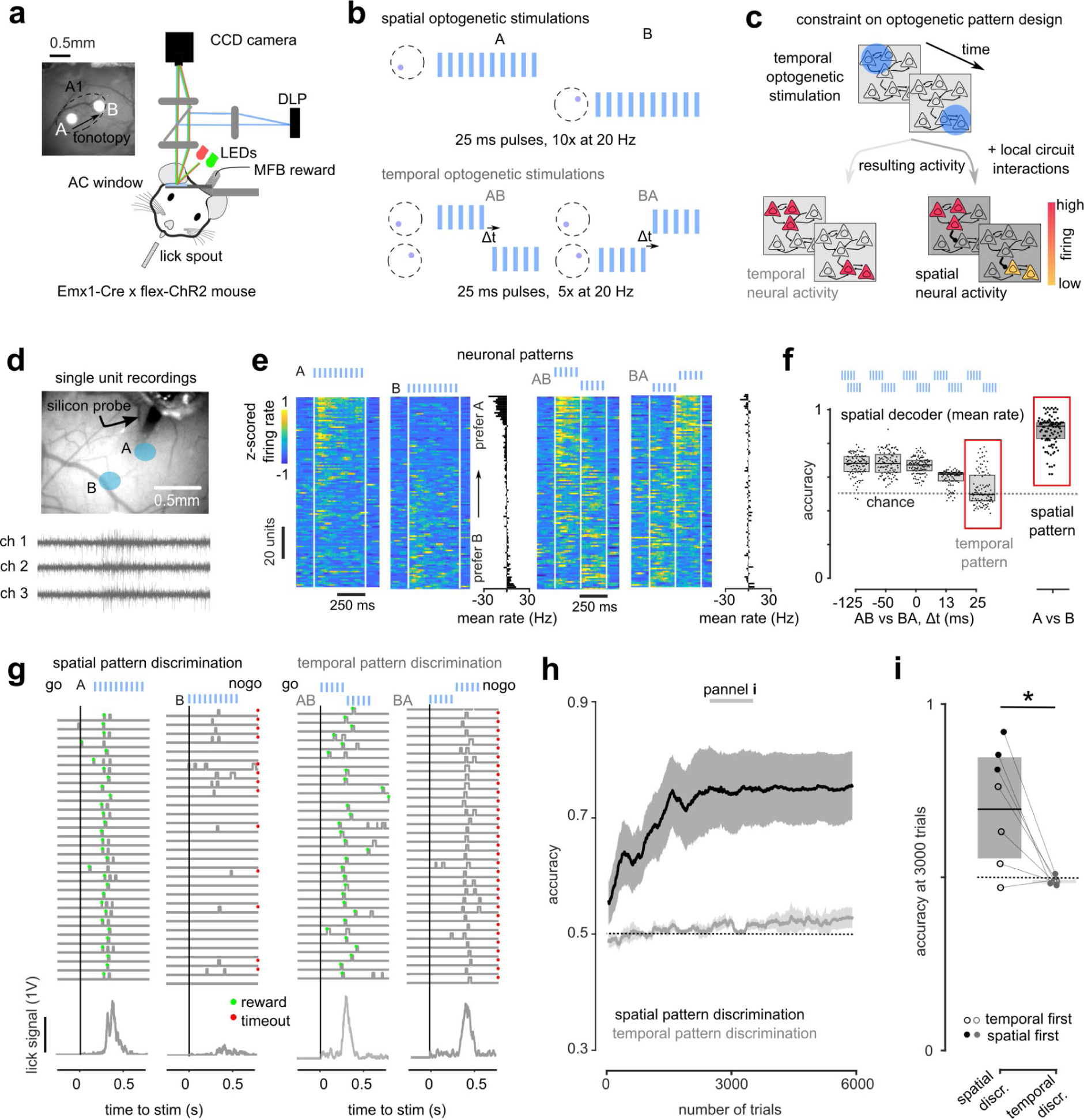
Sensory-motor learning requires spatial representations. **a.** Sketch of patterned optogenetic experiment in AC (MFB: medial forebrain bundle) and cranial window from an example mouse showing the location of the stimulation spots in the tonotopic axis of the primary auditory field. **b.** Sketch of the optogenetic stimulation time courses for each discrimination task. **c.** Sketch illustrating the conversion of a temporal code into a spatial code in the cortical network. **d.** AC window with 64 channel silicone probe inserted via a hole in the coverglass (top right) to record single unit responses to light patterns used in the behavioural task and illustrative data from 3 channels. **e.** Z-scored responses of 321 neurons to A, B, AB and BA stimulations, ordered by preference for A vs B stimulation and difference in average firing rate between A and B and between AB and BA stimulations. **f.** Accuracy of a neural decoder trained to discriminate the temporal and spatial optogenetic patterns based only on spatial information, i.e. time-averaged firing rates of each neuron (n=321 units, bootstrap over units, p-value of accuracy vs chance level of 0.5: 0.01, 0.01, 0.01, 0.43). **g.** Sample lick traces (top) and mean lick signal (bottom) for Go and NoGo trials in the rate-coded (left) and temporal-coded (right) discrimination tasks. **h.** Learning curves for all mice performing each task (n=7, error bars are sem). **i.** Accuracy at 3000 trials for all mice. (paired Wilcoxon test, p= 0.032, signed rank value = 27, n=7).

### A spatial code is necessary for learning

Mice were trained to discriminate the spatial (A vs B) and the temporal (A-B vs B-A) pairs of patterns in two Go/Nogo tasks. The tasks consisted in licking within a 1.5s opportunity window after the Go pattern onset to get a reward provided by medial forebrain bundle stimulation, which leads to identical learning rates as water rewards^40^. Licking for the NoGo pattern was punished by a timeout (**Fig. 1g**). Mice were trained on both tasks, counterbalancing task order. Consistent with previous results^29^, mice rapidly learned to discriminate in the spatial task, reaching 70% accuracy within 69+/-385 trials (**Fig. 1g-i, Extended Data Fig. 1b**). By contrast, learning was extremely slow and inefficient for the temporal task. After 3000 training trials, none of the mice could discriminate the temporal patterns, while most of them could discriminate the spatial patterns (**Fig. 1g-i**). Pushing training to even higher trial numbers, only two of seven mice reached slightly above chance levels for the temporal task (**Extended Data Fig. 1e**). This shows that activity patterns of cortical activity which contain temporal but no spatial information are hard to access for downstream learning processes for sensory-motor associations, while spatial patterns are easily exploited.

### Cortex elaborates a spatial code

We reasoned that this important constraint on AC output activity must have consequences for sound encoding across the auditory system. Many sounds differ only by temporal cues. For example a word and its time-reversed rendition are perceptually distinct in humans^5^. Rodents, including mice, are also able to discriminate between two sounds which mirror each other in time, an ability that depends on AC^41,42^. We therefore hypothesised that the inefficiency of circuits downstream of the cortex in learning only from temporal patterns constrains the auditory system to re-encode auditory temporal cues as spatial activity patterns. To test this hypothesis, we performed large-scale recordings in the awake mouse in three successive regions of the auditory system: the inferior colliculus (IC), the auditory thalamus (TH) and the AC (**Fig. 2a, Extended Data Table 1**), combined with auditory nerve (AN) responses simulated with a detailed biophysical model. In each region, we measured the responses to a set of 140 sounds, mainly of 500 ms duration, which covered simple, spectral and temporal features (**Fig. 2b, Extended Data Table 2**). In the IC, we used silicon probe electrophysiology to record 563 single units in the primary IC (central nucleus of IC) and 2-photon calcium imaging to record 13.132 ROIs from the more superficial secondary IC (dorsal cortex of IC). We also imaged 39.191 TH axonal boutons spread throughout AC and recorded 498 single units directly in TH. Finally, we imaged 60.822 ROIS throughout all subregions of the AC down to layer V. Calcium signals were linearly deconvolved ^24^, providing a temporal resolution of ∼150 ms sufficient to follow slow temporal patterns produced by our 500ms sounds. Full details of the dataset are provided in the **Supplementary Information** and **Extended Data Figs. 3 and 4**.

**Fig. 2.**
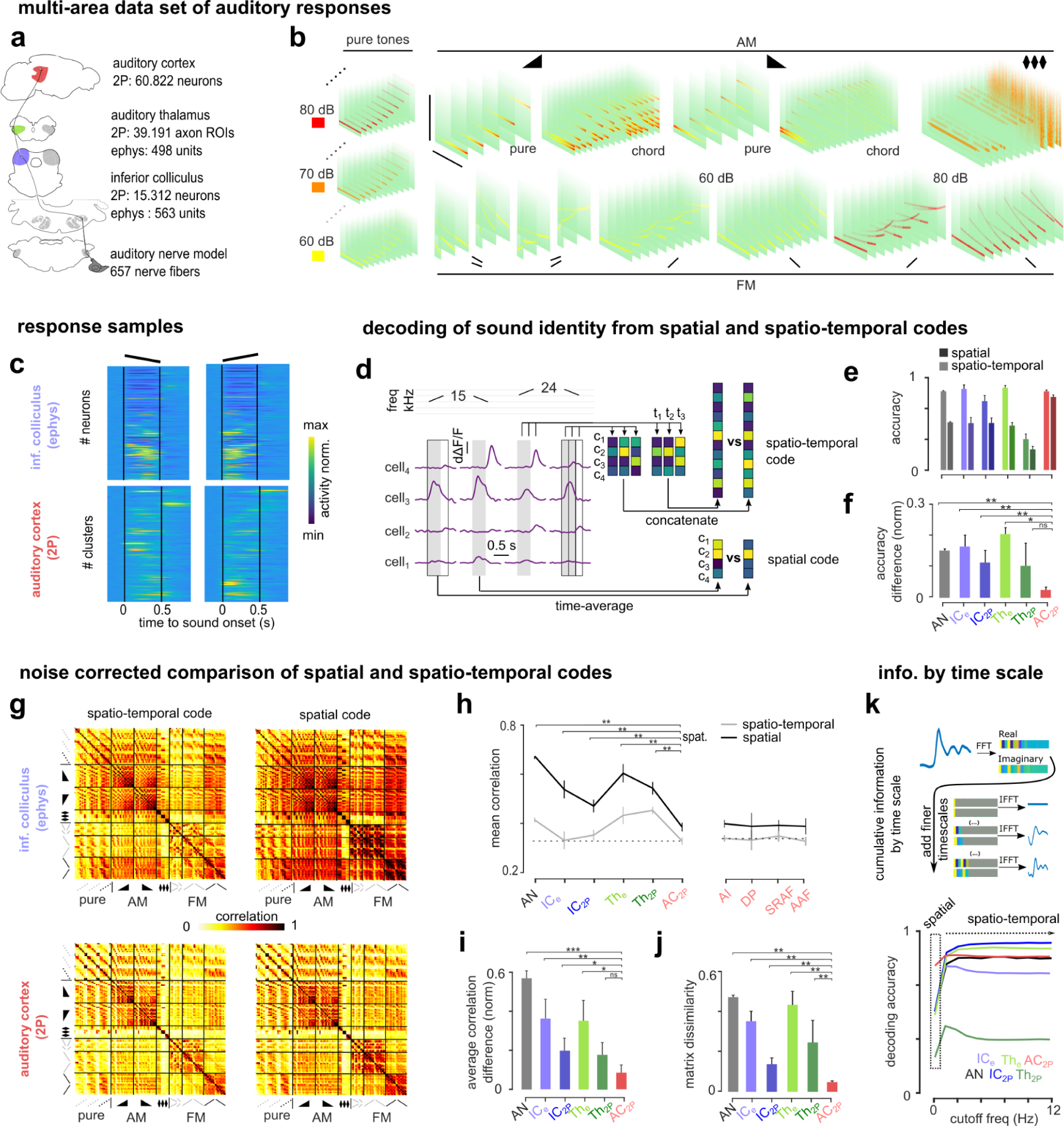
A spatial code for temporal cues emerges in the auditory cortex. **a.** Sketch of the auditory system and sample sizes at each level. **b.** Spectrograms of the sound set. **c.** Sample responses to up and down frequency sweeps from IC and AC neurons ordered by response amplitude. **d.** Responses of 4 AC neurons to different up and down frequency sweeps illustrating how spatio-temporal and spatial codes are extracted. **e-f.** Mean sound decoding accuracy for spatial-temporal and spatial codes in each area (**e**) and normalised difference between the two (**f**). (p-value for 100 bootstraps, error bars are S.D). **g.** Noise-corrected RSA matrices for all sound pairs for spatio-temporal (left) or spatial (right) codes in IC and AC. **h.** Mean noise-corrected correlation by area. (p-value for 100 bootstraps comparing rate correlation of each region to AC, error bars are bootstrapped S.D). **i.** Normalised difference between mean noise-corrected correlation for spatio-temporal and spatial codes. (p-value for 100 bootstraps, error bars are S.D). **j.** Noise-corrected dissimilarity between RSA matrix structure of spatio-temporal and spatial codes. (p-value for 100 bootstraps, error bars are S.D). **k.** (top) Sketch illustrating the decomposition of population responses by timescale and the concatenation of successive Fourier coefficients to accumulate increasingly fine timescales. (bottom) Mean decoding accuracy based on cumulative Fourier coefficients of neural responses. Full statistics are reported in **Extended Data Table 3**.

Contrasting neural responses to a frequency sweep and its time-reversed rendition (**Fig. 2c**) provides qualitative insight into the transformation of temporal cue representations that we observed throughout the auditory system. In the IC, the two sounds are represented by spatio-temporal activity patterns that involve the same neurons and mirror each other in time (**Fig. 2c**). In the AC, the temporal symmetry is no longer apparent and each sound is instead encoded by spatio-temporal activity patterns involving different neurons. This suggests that in AC but not in IC, sounds that differ only temporally are encoded by activity patterns that also differ in the spatial domain (**Fig. 2c**). This would make temporal information necessary in IC but dispensable in AC to discriminate between these sounds. We quantified this using two types of population activity decoders. To exhaustively extract the information contained in activity patterns, we used a spatio-temporal decoder that classifies sound responses with the full temporal sequence of population vectors (**Fig. 2d**). To extract information contained exclusively in spatial patterns, we used a spatial decoder that classifies sound responses with the population vectors obtained after time-averaging neuronal activity over the sound response (**Fig. 2d**). In subcortical areas, the spatio-temporal decoder clearly outperformed the spatial decoder. By contrast, in the cortex, the spatio-temporal and spatial decoder accuracies reached almost the same level (**Fig. 2e-f**). This result holds when the numbers of neurons are matched across datasets (**Extended Data Fig 5a**). Improved spatial decoding accuracy in AC could either result from a change of the representation as suggested in **Fig. 2c**, or from a change in the signal-to-noise ratio which varies across datasets (**Extended Data Fig. 5b**). To rule out the latter possibility, we quantified the similarity between population vectors evoked by a given sound pair. We used a numerically and analytically validated noise-corrected version of the Pearson correlation^43^ (**Extended Data Fig. 5c**, see **Supplementary Information** for mathematical derivations) to estimate similarity across representations in absence of noise.

Resulting representational similarity analysis (RSA) matrices summarise the relations between sound representations based either on the spatial or on the spatio-temporal information (**Fig. 2g, Extended Data Fig. 6a**). We first observed that, subcortically, the mean similarity between representations of different sounds (mean of RSA matrix) is higher for the spatio-temporal than for the spatial code, indicating that temporal patterns help segregating sounds in distinct representations. However, this difference is very small in AC contrary to subcortical structures (**Fig. 2h,i**). In addition RSA matrices for spatial and spatio-temporal representations were very similar in AC, while they were much dissimilar subcortically (**Fig. 2j**). These results hold in all subfields of AC (**Fig. 2h**) and are robust to the number of neurons included in the analysis (**Extended Data Fig. 5d**). This together demonstrates that spatio-temporal and spatial representations of sounds in AC are very similar, explaining why decoders perform almost equally with both representations. Together, these results show that time-varying sounds are accurately represented by purely spatial patterns in the AC but not in subcortical structures. Interestingly, this transformation makes temporal sensory information available to learning mechanisms requiring a spatial representation, reconciling our optogenetic results with the ability to discriminate temporal cues in sounds.

### Spatial and temporal codes co-exist

We then tested if the convergence between spatio-temporal and spatial representations in AC is the consequence of a decrease in temporal resolution in the cortex^22,44^ or occurs without loss of temporal information. We decomposed neural population activity using Fourier analysis and measured classifier accuracy at each specific timescale (**Fig. 2k, Extended Data Fig. 5e**). This analysis showed that relevant temporal resolution is preserved on the time scales considered in our study. In all datasets, classifier accuracy is maximal at ∼1.5 Hz resolution (**Extended Data Fig. 5e**). Thus all datasets contain temporally structured neural activity that is sufficient to identify the relevant temporal cues in our sounds. Moreover, accumulating information from low to high temporal resolutions shows a saturation of classifier accuracy at around 3 Hz for all datasets (**Fig. 2k**). Therefore all temporal information needed to discriminate our sounds is available below 3 Hz, which is much lower than the putative 30 Hz cutoff for temporal resolution in AC ^22^. We observed sound-related information at fast timescales, in particular in electrophysiological recordings (**Extended Data Fig. 5e**) but it was redundant to information at slower time scales (**Fig. 2k**). Corroborating the spatial and spatio-temporal decoders (**Fig. 2e,f**), the time-averaged activity of neurons (0 Hz) reached a level similar to that of the full cumulative spatio-temporal information in AC but not subcortically (**Fig. 2k**). Therefore, in AC, almost all information, including that present in neural temporal patterns, is also accessible from the identity of active neurons, i.e. from a purely spatial code. Accordingly, each sound is represented by small sets of highly active, and highly specific neurons in AC as shown by the high population and lifetime sparseness (**Extended Data Fig. 5f,g**)^45^. Notably, this property evolved non-monotonically along the auditory system, with much less sparse representations in TH (**Extended Data Fig. 5f,g**), paralleling the increase in representation similarity in this area (**Fig. 2h**). In contrast to sparseness measures, the level of tuning to simple, individual temporal features (e.g. frequency modulation direction) was stable from IC to AC (**Extended Data Fig. 7**), consistent with previous reports^46,47^. This suggests that the cortical transformation of sound representations does not correspond to a sharpening of particular tuning properties but to the emergence of more complex tuning properties^48^.

### The spatial code sets learning speed

We next investigated in a computational model if known learning mechanisms downstream of cortex can capture the advantages of the cortical elaboration of a spatial code for temporal information. We used a feedforward neural network model which simulates discrimination learning in an auditory Go/NoGo task based on reinforcement learning principles^49^ (**Fig. 3a**). We upgraded this model by complementing its Hebbian synaptic learning rule with an eligibility trace mechanism^50,51^ parameterized with data from the striatum^52^, a structure receiving AC projections and implicated in sound discrimination learning^33^. The eligibility trace flags active synapses with a signal that decays over ∼1-3s. This mechanism allows even delayed post-synaptic activity driven by the reward to gate plasticity based on presynaptic input (**Fig. 3a**). However, when associated with Hebbian plasticity, the long decay of the eligibility trace averages out the precise timing of pre-and postsynaptic activity coincidences. Thus, when we trained the model to discriminate between the population responses to pairs of sounds taken from the AC, TH or IC datasets, we observed that the model learned from the spatial patterns but ignored the temporal patterns. This can be evidenced by plotting learning duration as a function of the noise-corrected similarity (correlation in RSA matrix) between discriminated representations, which shows that learning duration is much more correlated to the spatial than to the spatio-temporal similarity (**Fig. 3b)**. Moreover, learning duration rises in a steep and non-linear manner for high spatial representation similarity (**Fig. 3c**). This non-linear relationship matches the result of our optogenetic experiments (**Fig. 1**), in which temporal sequences with maximal spatial representation similarity yielded extremely slow learning. The model also provides an explanation for the long standing observation that discrimination of pure tones does not require AC and can be performed via subcortical sensory-motor projections, whereas discrimination of temporal cues requires AC^29,41,42,53^ (**Fig. 3d**). Simple sound pairs, such as pure tones differing enough in frequency (e.g. > 0.33 octave), have low spatial representation similarity at all stages of the auditory (e.g. correlation < 0.75, **Fig. 3e, Extended Data Fig. 6b-d**). For this range of low correlation values, our model shows that learning occurs quickly and the impact of representation similarity on learning speed is marginal (**Fig. 3c**). Hence, the model predicts similar learning speeds whether it is based on thalamic or cortical representations (**Fig. 3e**), as observed for pure tone discriminations with intact or ablated AC^41^. Contrariwise, sounds that differ only in their temporal structure, such as time-symmetric frequency modulations, have spatial representations that are highly correlated subcortically (>0.9, **Fig. 3f**) and clearly less in the cortex (0.74, **Fig. 3f**, similar results for other temporal cues, **Extended Data Fig. 6e-g**). Based on these values, our model predicts a ∼3-fold decrease in learning duration with cortical compared to thalamic representations (**Fig. 3f**). This is in line with the observation that pre-training AC ablation severely prolongs discrimination learning for time-reversed frequency sweeps^41^. Also, if one postulates that learning speed determines which auditory system stage is recruited for solving a sound discrimination task, the relationship between spatial representation and learning duration in our model can explain the strong impact of post-training AC inactivation for discrimination of temporal cues, but not for pure tones^29,42,53^. This together indicates that the properties of learning rules in the striatum can quantitatively explain the necessity of an upstream spatial code for sensory-motor learning and recapitulate effects of causal manipulations of AC on auditory discrimination learning.

**Fig. 3.**
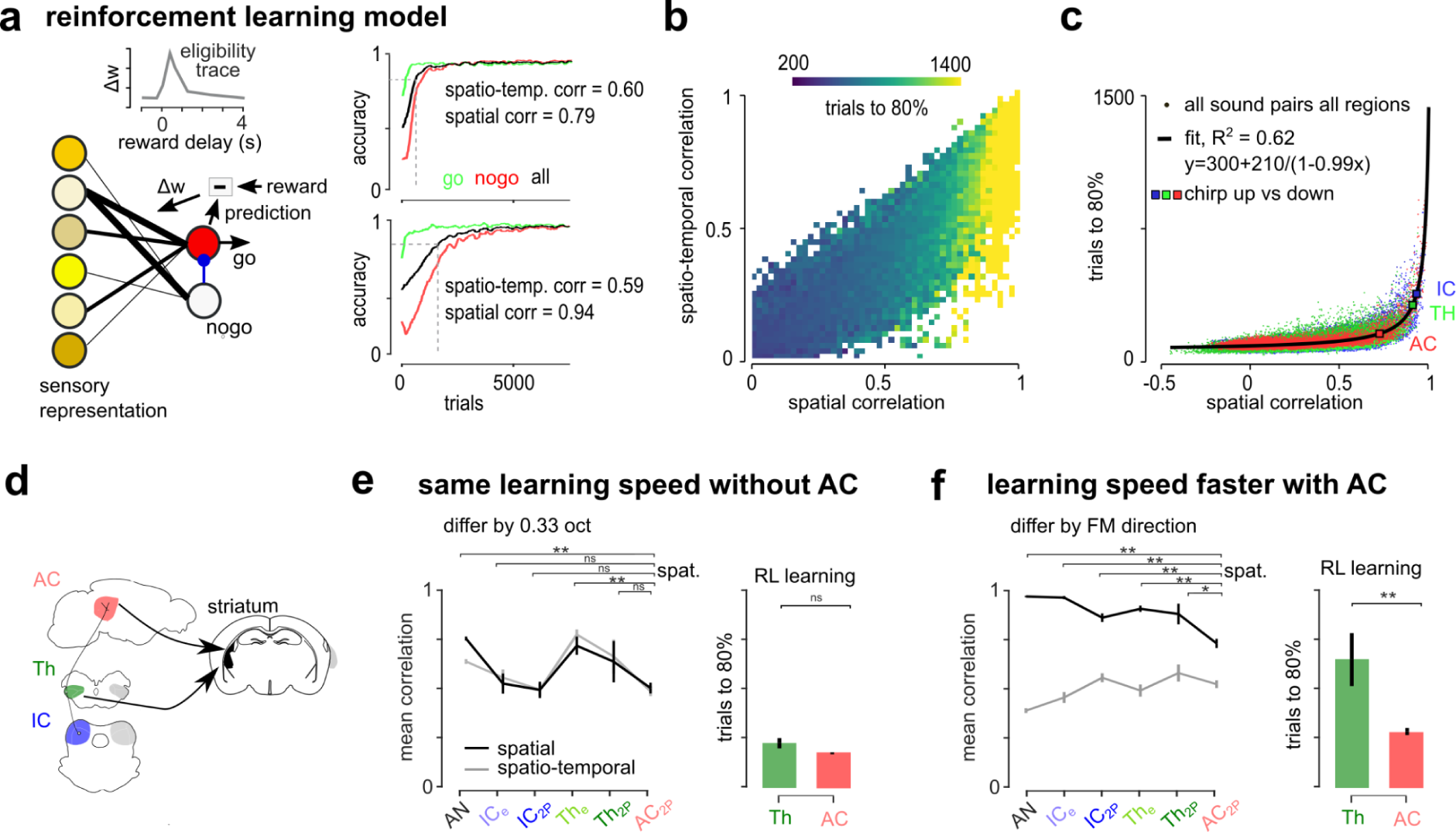
A spatial code is necessary for reinforcement learning with a bio-inspired eligibility trace mechanism. **a.** Sketch of the reinforcement learning model (bottom left), eligibility trace dynamics (top left) and example learning curves for two recorded representations that have similar spatio-temporal correlations but different spatial correlations. **b.** Heatmap of the number of trials needed to reach 80% accuracy at discriminating between a pair of sounds as a function of the correlations of their spatio-temporal and spatial representations. The colour map indicates learning duration averaged over all pairs of representations for all brain regions. **c.** Number of trials to 80% accuracy as a function of the correlations of their spatial representations. Large square dots show the mean correlation and learning time for time-symmetric frequency sweeps in IC, TH and AC and the black line shows the fit to data. **d.** Sketch showing the thalamic and cortical pathways for auditory learning. **e.** Mean noise-corrected correlation between representations of sound pairs differing only by frequency (0.33 octave difference) and predicted duration for learning a pure tone discrimination task based on thalamic (average of THe and TH2P) and cortical representations. **f.** Mean noise-corrected correlation between representations of sound pairs differing only by the direction of the frequency sweep and predicted duration for learning to discriminate the two frequency sweep directions based on thalamic (average of THe and TH2P) and cortical representations. Full statistics are reported in **Extended Data Table 3**.

### Spatial codes emerge from categorisation

The emergence of a spatial code for temporal information in the cortex may more generally reflect computations related to the resolution of common perceptual tasks such as stimulus identification and categorization. To explore this theoretically, we analysed representations in convolutional neural networks (CNNs) trained for different sound processing tasks (**Fig. 4**). A first network was trained at categorising key features of the stimuli presented to our mice: the frequency and intensity range, and the type of frequency and amplitude modulations present in the sounds (**Fig. 4a, Extended Data Fig. 8a**). This network generated a spatial code for temporal cues in its deep layers after training, as shown by the convergence of spatial and spatio-temporal similarity (**Fig. 4a, Extended Data Fig. 8b**). Like typical CNNs, our network implemented pooling mechanisms which increase the size of sensory receptive fields and shrink the temporal and spatial dimensions in deeper layers^54^. To rule out that the convergence between spatio-temporal and spatial codes is due to this temporal shrinking, we trained a second CNN without pooling over the temporal dimension (**Fig. 4b**). We observed that temporal shrinking accelerates learning but is not necessary for the emergence of a spatial code for temporal cues in the CNNs (**Fig. 4b, Extended Data Fig. 8a**). This network displayed properties similar to the auditory system (compare **Fig. 4c,d** and **Fig. 2f,j**) and in particular the spatial code for temporal cues in deep CNN layers did not involve a decreased temporal resolution of the representation (compare **Fig. 4e, Extended Data Fig. 8c** and **Fig. 2k, Extended Data Fig. 5e**). These results extended to a previously published CNN trained to classify words and musical styles^55^ (**Extended DataFig. 8d**) and to networks trained to perform single sound identification in noise (**Extended DataFig. 8e**). However, when we trained another CNN network, with an auto-encoder architecture, to compress and denoise sound representations without assigning specific labels to sounds, we did not observe the emergence of a spatial code for temporal cues (**Fig. 4f, Extended Data Fig. 8f,g**). Therefore, CNN models support the view that the emergence of spatial representations for temporal cues in the cortex is driven by the computational constraints of classifying sounds into perceptual objects assigned with meaning.

**Fig. 4.**
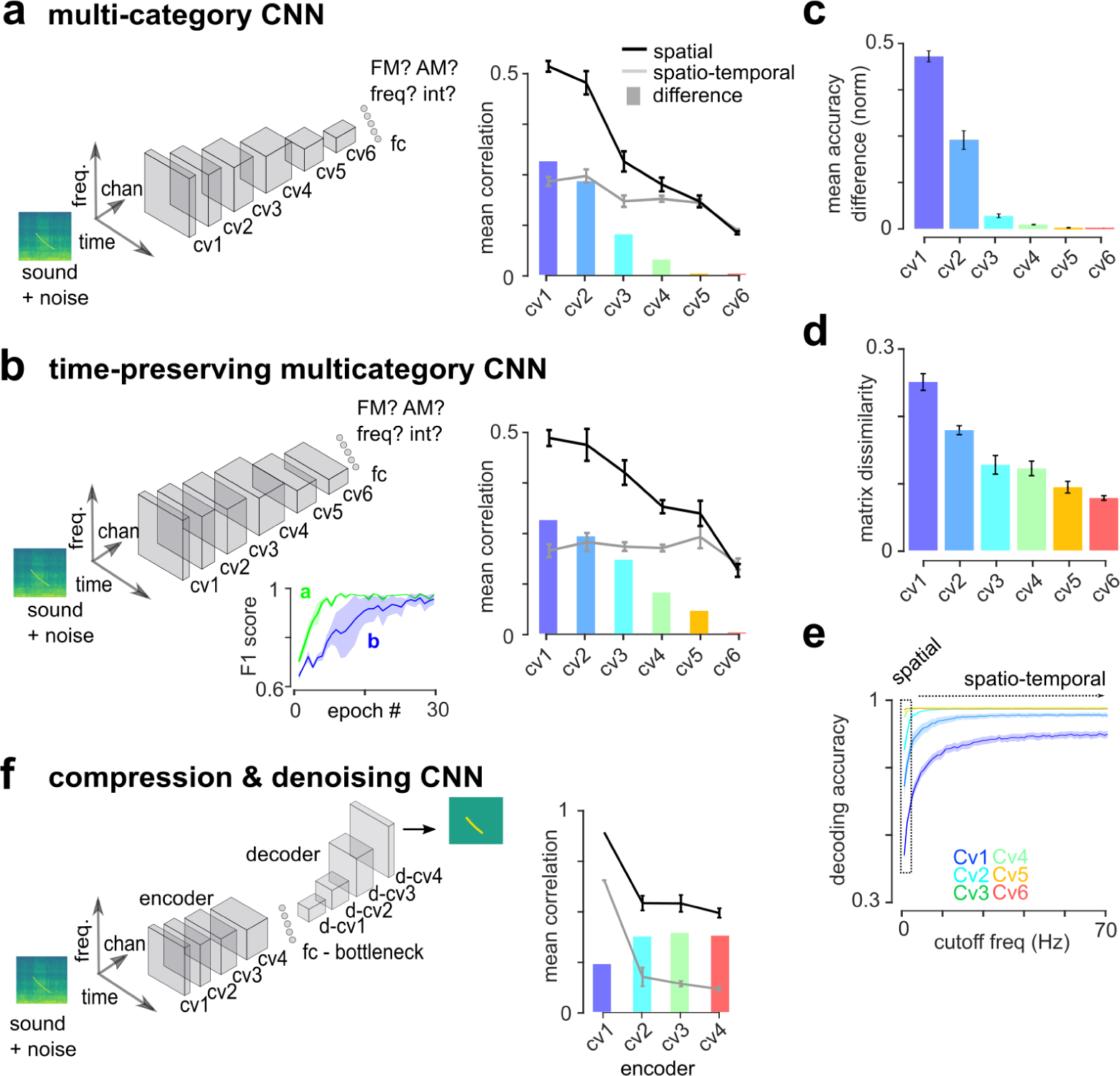
Categorization deep networks implement a spatial code for temporal cues in deeper layers. **a-b.** (Left) Schematic of CNN architectures and target categories. (Right) Mean response correlations for the spatial and spatio-temporal codes from RSA matrices constructed with the set of 140 sounds presented to mice (line) and difference between the two codes (bars). **a.** Multi-category CNN (n=8 networks). **b.** Multi-category CNN without shrinking of the temporal dimension (n=8 networks). Inset shows learning curves from training epochs for networks in A and B. **c-e.** All graphs refer to the categorization CNN without temporal pooling and reproduce analysis shown in Fig. 2 for neural data. **c.** Normalised difference between mean sound decoding accuracy for spatio-temporal and spatial codes. (error bars are sem over trained networks). **d.** Noise-corrected dissimilarity between RSA matrix structure of spatial and spatio-temporal codes. **e.** Mean decoding accuracy based on cumulative Fourier coefficients of neural responses. **f.** Autoencoder CNN performing sound compression and denoising through a 20-unit bottleneck. (cv : convolution block, d-cv : deconvolution block - see methods for architecture details).

## Discussion

Our results highlight the spatial encoding of temporal sound cues as an important function of the sensory cortical network. This is in line with the proposed role of AC in the encoding of auditory objects^56^ such as phonemes, vocalisations or musical notes and previous observations of spatial representations for speech or natural sounds in the AC of humans and animals^10,55,57^. Our results demonstrate that this spatial encoding is necessary for rapid learning of sensory-motor associations. Based on a reduced model of sensory discrimination learning, we propose that this constraint arises from time-averaging properties of plasticity rules implemented in associative centres such as the striatum or the amygdala. These brain regions link information from a wide range of sensory areas to behavioural responses and environmental outcomes that have their own, unrelated temporal structure. Our results suggest that the challenge of associating the distinct temporalities of sensory signals and motor responses is resolved via the use of spatial representations for temporal cues.

Several computational models^58–60^ and experimental findings^61,62^ have identified plasticity mechanisms by which temporal sequences can be associated to a specific neuronal output, by combining spike-timing-dependent synaptic plasticity (STDP)^63^ and neuronal integration mechanisms. Several factors could explain why such mechanisms are not efficiently recruited in auditory sensory-motor learning. First, many auditory objects, like our optogenetic stimuli, evolve on timescales of hundreds of milliseconds that are not suited for the short timescales of STDP^64^. Second, different models indicate that under irregular spike train statistics as observed *in vivo*, STDP rules behave as standard Hebbian rules^60,65,66^. While temporally precise sound responses in AC were often reported under anaesthesia, more irregular spike trains are observed in the awake state^67^. Likewise, our mild optogenetic stimulations calibrated to yield realistic firing rates (**Extended Data Fig. 1d**) are likely too weak to overcome cortical noise^39^ and generate high temporal precision (**Fig. 1e**). Finally, eligibility traces gate plasticity based on neuromodulatory feedback which, in the case of striatal dopamine, can occur up to 3s after synaptic activity. This slow timescale of integration averages out the precise timing of pre- and postsynaptic activity coincidences and could explain why fine temporal information is inaccessible to plasticity mechanisms that are driven by delayed environmental feedback.

A previous study showed that rats can discriminate short time intervals between two electrical stimulations at two different AC locations^36^. We expect these results to reflect both the acute precision of electrical stimulation and the conversion of fast temporal information into a spatial code via synaptic interactions in the cortical networks (**Extended Data Fig. 2**), as we observed when decreasing the time interval in our paradigm (**Fig. 1f**). Network interactions in AC may underlie the transfer of temporal information received from thalamus into the spatial domain (**Fig. 2**). CNNs are also based on local computations that iteratively detect spectrotemporal features and recapitulate the emergence of the spatial code (**Fig. 4**). Hence, simple local computations whose implementation in the local circuit motifs of the auditory system remain to be defined, are sufficient to make temporal information accessible through a spatial code in AC. Our study also revealed two intriguing aspects of sound information processing in the auditory system. First, contrary to what is observed in CNNs (**Fig. 4**), representations from IC to AC are transformed non-monotonically with denser, more correlated representations in TH compared to AC and IC (**Fig. 2**). Second, it is remarkable that neural temporal information is largely preserved in the AC despite the accessibility of temporal information through a spatial code (**Fig. 2**). This coexistence of temporal and spatial coding schemes could serve to combine object-like representations, useful for categorical decisions, with an explicit representation of the temporal details that are also perceived together with the object.

## Methods

### Subjects and authorizations

All mice used for imaging and electrophysiology were 6 to 14 weeks old male and female C57Bl6J mice that had not undergone any other procedures. For optogenetic stimulation, we used Emx1-IRES-Cre (Jax #005628) crossed with Ai27 (Jax #012567) mice. Mice were group-housed (2–6 per cage) before and after surgery, had *ad libitum* access to food and water and enrichment (running wheel, cotton bedding and wooden logs) and were maintained on a 12-hour light-dark cycle in controlled humidity and temperature conditions (21-23°C, 45-55% humidity). All experiments were performed during the light phase. All experimental and surgical procedures were carried out in accordance with the French Ethical Committee the French Ethical Committees #59 and #89 (authorizations APAFIS#9714-2018011108392486 v2 and APAFIS#27040-2020090316536717 v1).

### Surgery

Mice were injected with buprenorphine (Vétergesic, 0,05-0,1 mg/kg) 30 min prior to surgery. Surgical procedures were carried out using either intraperitoneal ketamine (Ketasol) and medetomidine (Domitor) which was antagonised with atipamezole (Antisedan, Orion pharma) at the end of the surgery) or 3% isoflurane delivered via a mask. After induction, mice were kept on a thermal blanket during the whole procedure and their eyes were protected with Ocrygel (TVM Lab). Lidocaine was injected under the skin of the skull 5 minutes prior to incision.

For calcium imaging, craniotomies of 3 (IC) or 5 (AC) mm were performed above the IC or the AC. Injections of 150nL of AAV1.Syn.GCaMP6s.WPRE (Vector Core, Philadelphia, PA; 10^13 viral particles per ml; used pure for TH and diluted 30x for AC and IC) were made at 30 nL/min with pulled glass pipettes at a depth of 500µm and spaced every 500 µm to cover the a large surface of the IC or AC. The craniotomy was sealed with a circular glass coverslip. The coverslip and head post were fixed to the skull using cyanolit glue and dental cement (Ortho-Jet, Lang).

For electrophysiology recordings, the skull above the IC or above the cortex dorsal to the TH was exposed for ulterior craniotomy. A well was formed around it using dental cement in order to retain saline solution during recordings and the head post was fixed to the skull using cyanolit glue and dental cement. To protect the skull, the well was filled with a waterproof silicone elastomer (Kwikcast, WPI) that could be removed prior to recording. The head post was fixed to the skull using cyanolit glue and dental cement (Ortho-Jet, Lang).

For patterned optogenetic stimulation of the cortex, a cranial window was placed above the AC as for calcium imaging but without viral injection. For MFB stimulation, a bipolar stimulation electrode (60-µm-diameter twisted stainless steel, PlasticsOne) was implanted using stereotaxic coordinates (AP −1.4, ML +1.2, DV +4.8). It was then fixed along with the headplate to the skull using dental cement (Ortho-Jet, Lang).

After surgery, mice received a subcutaneous injection of 30% glucose and metacam (1 mg/kg). Mice were subsequently housed for one week with metacam delivered via drinking water or dietgel (ClearH20). Mice were given one week to recover from surgery without any manipulation. Then, for four days before recording, mice were habituated to head restraint for increasing periods of time (30 min - 2 hours). For electrophysiological experiments, the day before recording animals were briefly anaesthetised using isoflurane anaesthesia (2%) in order to perform craniotomy and durectomy for electrode descent.

### Two photon calcium imaging in the awake mouse

Imaging was performed using a two-photon microscope (Femtonics, Budapest, Hungary) equipped with an 8kHz resonant scanner combined with a pulsed laser (MaiTai-DS, SpectraPhysics, Santa Clara, CA) set at 900 nm. We used a 10x Olympus objective (XLPLN10XSVMP), which provided a field of view of up to 1×1 mm. For AC, a 1×1mm field of view was used. For IC, the field of view was adjusted to the size of the structure (∼0.5×0.5 mm). For thalamic axons, the field of view was reduced to 0.22×0.22 mm. Images were acquired at 31.5 Hz.

### Electrophysiology in the awake mouse

Electrophysiology was performed using Neuronexus probes : (1×32 linear probe for IC and 4*8 comb for TH). For track reconstruction, the electrodes were dipped in diI, diO or diD (Vybrant™ Multicolor Cell-Labelling Kit, Thermofisher) prior to recording and allowed to dry at least 15 min before insertion. Recordings were performed using warmed saline filling the cyanolit glue well and in contact with the reference electrode. After each recording the well was amply flushed and then refilled with Kwickast. A maximum of three recordings were performed per site. Data was sampled at 20kHz using an Intan RHD2000 amplifier board.

For recordings during optogenetic stimulation, a small hole was drilled in the coverglass and a 1×64 linear probe was inserted into the stimulated region. During these recordings, optogenetic stimuli and sounds were presented randomly. Recordings were performed using warmed saline filling the cyanolit glue well and with a reference electrode chronically implanted into the brain. After each recording the well was amply flushed and then refilled with Kwickast.

### Sound delivery

Sounds were generated with Matlab (The Mathworks, Natick, MA) and were delivered at 192 kHz with a NI-PCI-6221 card (National Instruments) driven by the software Elphy (G. Sadoc, UNIC, France) and feeding an amplified free-field loudspeaker (SA1 and MF1-S, Tucker-Davis Technologies, Alachua, FL) positioned 15 to 20 cm from the mouse ear. Sound intensity was cosine-ramped over 10 ms at the onset and offset to avoid spectral splatter. The head fixed mouse was isolated from external noise sources by sound-proof boxes (custom-made by Femtonics, Budapest, Hungary or Decibel France, Miribel, France) providing 30 dB attenuation above 1 kHz. Sounds were calibrated in intensity at the location of the mouse ear using a probe microphone (Bruel & Kjaer, type 4939-L-002). For two-photon calcium imaging, the resonant scanner generated a harmonic background noise at 8 kHz (intensity at the mouse ear, 45 dB SPL).

During a recording session, each of the 140 sounds (sketched in **Fig. 2b**) was presented 15 times in random order. In order to be compatible with 2-photon image acquisition, sounds were presented in 120 blocks of 32s each, interleaved by a 15s pause in a 94 min protocol. The list of all sound parameters can be found in **Extended Data Table 2**.

### Intrinsic optical imaging recordings in anaesthetised mice

Intrinsic imaging was performed to localise AC in mice under light isoflurane anaesthesia (1% delivered with SomnoSuite, Kent Scientific) on a thermal blanket. Images were acquired at 20Hz using a 50mm objective (1.2 NA, NIKKOR, Nikon) with a CCDcamera (GC651MP, Smartek Vision) equipped with a 50 mm objective (Fujinon, HF50HA-1B, Fujifilm) through the cranial window implanted 1-2 weeks before the experiment (4-pixel binning, field of view between 3.7 x 2.8 mm or 164 x 124 pixels at 5.58 mm/pixel). Signals were obtained under 780 nm LED illumination (M780D2, Thorlabs). Images of the vasculature over the same field of view were taken under 530 nm LED illumination (NSPG310B, Conrad). Sequences of short pure tones at 80 dB SPL were repeated for 2 s every 30 s with 10 trials per sound. Acquisition was triggered and synchronised using a custom-made GUI in MATLAB. For each sound, we computed baseline and response images, 3 s before and 3 s after sound onset, respectively. The change in light reflectance ΔR/R was calculated for each repetition of each sound frequency (4, 8, 16, 32 kHz, white noise) as the difference between the baseline and response image and was then averaged across all repetitions of a given tone frequency. Response images were smoothed applying a 2D Gaussian filter (sd = 3 pixels). Auditory cortex activity appeared as regions with reduced light reflectance changing with frequency, revealing the tonotopic maps of its different subfields. To align intrinsic imaging responses across different animals, the 4 kHz response was used as a functional landmark. The spatial locations of maximal amplitude responses in the 4 kHz response map for the A1, A2 and AAF (three points) was extracted for each mouse and a Euclidean transformation matrix was calculated by minimising the sum of squared deviations (RMSD) for the distance between the three landmarks across mice. This procedure yielded a matrix of rotation and translation for each mouse that was applied to compute intrinsic imaging responses averaged across a population of mice.

### Histology and immunostainings

In order to extract the brain for histology, mice were deeply anaesthetised using a ketamine-medetomidine mixture and perfused intracardially with 4% buffered paraformaldehyde fixative. The brains were dissected and left in paraformaldehyde overnight and then sliced into fifty micrometre sections using a vibratome. Slices were either stained with cytochrome oxidase or directly mounted using a mounting medium with DAPI. Analysis of the fluorescence band diI, diO or diD allowed isolating up to 3 tracks per mouse for electrophysiological experiments.

For Vglut2 immunostainings, after fixation, tissues were rinsed in PBS and blocked in Tris-Buffered Saline (TBS) supplemented with 5 % (vol/vol) Normal Donkey Serum (Jackson Immunoresearch) and 0.3 % (wg/vol) Triton X-100. Then, sections were incubated for 48h at 4°C while rocking with a primary antibody: guinea pig anti-Vglut2 (1:500, Synaptic Systems #135404), followed by a 4 h incubation with a secondary donkey anti-guinea pig IgG [F(ab’)2fragments] (1:500, Jackson ImmunoResearch #706606148). Tissues were rinsed and mounted using Prolong diamond antifade (Life Technologies). Pictures of the brain sections were taken with LSM 900 confocal microscope (Zeiss Microsystems) using 20x objective, whereas the magnified view of the thalamocortical boutons was obtained with Airyscan acquisition and 63x objective.

The labelled boutons (GCaMP alone in green; GCaMP with Vglut2 in yellow) were counted manually using ZEISS ZEN 2 microscope software in 12 sample regions selected within layer 1 AC in 3 different Airyscan images. The number of boutons was then calculated per volume tissue.

### Behavioural discrimination of patterned optogenetic stimuli

For patterned optogenetic activation in the mouse AC, we used a video projector (DLP LightCrafter, Texas Instruments) powered by a blue LED (centre wavelength 460 nm). To project a two-dimensional image onto the AC surface. The image of the micromirror chip was collimated through a 150 mm cylindrical lens (Thorlabs, diameter: 2 inches) and focused through a 50 mm objective (NIKKOR, Nikon). Light collected by the objective passes through a dichroic beam splitter (long pass, > 640nm, FF640-FDi01, Semrock) and is collected by a CCD camera (GC651MP, Smartek Vision) equipped with a 50 mm objective (Fujinon, HF50HA-1B, Fujifilm).

The behavioural task aimed to teach mice to discriminate between two optogenetically induced patterns of activity in AC. The reinforcement used for the task used medial forebrain bundle (MFB) stimulation in non-deprived mice. This protocol leads to similar learning speed, motor response timing and psychometric measurements as water rewards in deprived animals^40^. In the “spatial discrimination task”, the two stimuli were composed of 500 ms illumination of 300 µm diameter spots placed at two different locations of AC. In the “temporal discrimination task”, the two stimuli were composed of a succession of two 250 ms illuminations of 300 µm diameter spots at different locations in the cortex in one order (AB) or in the reversed order (BA). All light stimuli were temporally modulated at 20 Hz (25 ms ON, 25 ms OFF). To prevent visual perception of the optogenetic stimuli a constant and strong background illumination provided by a white LED lamp was used and a cache was placed in front and close to the eyes to limit visual inputs. Mice were trained on both tasks. 4 mice were first trained on the temporal, and then on the spatial task. 3 mice were first trained on the spatial and then on the temporal task. The spots used in the task they first learnt were positioned at the two extremes of the tonotopic axis of A1. In order to minimise interference between the two subsequent tasks, the spots in the second task were positioned at two different locations, on the axis orthogonal to the tonotopic axis of A1 keeping the inter-spot distance equal. In both cases, spot position was adjusted to avoid placing them above major vessels which could lead to reduced illumination of neurons (**Extended Data Fig. 1a,b**). Alignment of optogenetic stimulus locations across days was done using blood vessel patterns at the surface of the brain manually aligned to a reference blood vessel image taken at the beginning of the experiment.

Behavioural experiments were monitored and controlled using a custom Matlab software controlling an input-output board (PCIe-6351, National Instruments) and the images delivered by the video projector. Mice performed behaviour for one hour per day. During the entire behavioural training period, food and water were available *ad libitum* as rewards were provided through the stimulation of the medial forebrain bundle (MFB).

MFB stimulation was delivered via a pulse train generator (PulsePal V2, Sanworks) that produced 2ms biphasic pulses at 50Hz for 100ms at a voltage calibrated for each individual mouse to the minimal level that evoked sustained responding, using the protocol in ^40^. The stimulation was controlled with a solenoid valve (LVM10R1-6B-1-Q, SMC). A voltage of 5V was applied through an electric circuit joining the lick tube and an aluminium foil on which the mouse was sitting. Lick events could be monitored by measuring the voltage across a series resistor in this circuit.

Training was broken down into three phases. (i) Lick training: On the first day, mice were presented with the lick tube and any licking was rewarded with immediate MFB stimulation. Mice generally began licking at high rates after 1-2 minutes and the session was continued until mice reliably collected around 300 rewards. (ii) Go training: On the following day, Go trials were presented with 80% probability, while the remaining trials were blank trials (no stimulus). A trial consisted of a random inter-trial interval (ITI : 0.5 to 1 s), a random ‘no lick’ period (duration adjusted, see below) and a fixed response window of 1.5 s. The first lick occuring during the response window on a Go trial was scored as a ‘hit’ and triggered immediate MFB stimulation. During initial go training the ‘no lick’ period was between 2 and 5 s in order to discourage non-specific licking. When mice achieved >80% accuracy for the Go stimulus, a final Go session was performed during which a cache was placed over the window to verify that animals were not licking to remnant visual cues from the video projector (Extended Data **Fig. 1c,d**). On this day and for subsequent Go/NoGo sessions, the no lick period was shortened to 1.5 to 3 s in order to obtain more trials per session. (iii) Go/NoGo training: After Go training, the second stimulus (NoGo) was introduced. During presentation of the NoGo sound, the absence of licking for the full response window was scored as a ‘correct rejection’ (CR) and the next trial immediately followed. Any licking during NoGo trials was scored as a ‘false alarm’ (FA), no stimulation was given, and the animal was punished with a random time-out period between 5 and 7 s. Each session contained 45% Go stimuli, 45% NoGo stimuli and 10% blank stimuli. Note that the Go training was used to ensure high motivation of the animal during the Go/Nogo training by establishing an association between the optogenetic stimulus and the reward. For the time-independent task, this association was generalised to the NoGo stimulus, as seen through very high false alarm rates at the beginning of the Go/NoGo training (e.g. **Extended Data Fig. 1b**). This indicates that faster learning for the time-independent task is not due to an absence of generalisation between the Go and NoGo stimulus when transitioning from the Go to the Go/NoGo training phases.

Learning curves were obtained by calculating the fraction of correct responses over blocks of 150 trials. Discrimination performance over one session was calculated as (hits + correct rejections)/total trials.

### Data pre-processing

For calcium imaging, regions of interest corresponding to putative neurons (AC and IC) or axons and boutons (TH) were identified by using Autocell ^24^ (https://github.com/thomasdeneux/Autocell). Briefly, each frame of the recording was corrected for horizontal motion using rigid body registration.This step was visually controlled and all sessions with visible z motion were eliminated. A hierarchical clustering algorithm, based on pixel covariance over time, agglomerated pixels up to a user-selected number of clusters corresponding to regions of the size of neurons of axons. Clusters were automatically filtered according to size and shape criteria. This step was controlled by a detailed visual inspection of selected regions of interest (ROIs) during which ROIs without visually identifiable cell body shape were discarded.

For each region of interest, the mean fluorescence signal F(t) was extracted together with the local neuropil signal F_np_(t). Then 70% of the neuropil signal was subtracted from the neuron signal to limit neuropil contamination. Baseline fluorescence F_0_ was calculated with a sliding window computing the 3rd percentile of a Gaussian-filtered trace over the imaging blocks. Fluorescence variations were then computed as f(t) = ΔF/F = (F(t) - F_0_)/F_0_. An estimate of firing rate variations r(t) was then obtained by linear temporal deconvolution of f(t): r(t) = f’(t) + f(t)/τ, f’(t) being the first derivative of f(t) and τ = 2s, the estimated decay of the GCAMP6s fluorescent transients. This simple method efficiently corrects the strong discrepancy between fluorescence and firing rate time courses due to the slow decay of spike-triggered calcium transients. It does not correct for the rise time of GCAMP6s, leading to remnant low pass filtering of the firing rate estimate and a delay of ∼100ms between the firing rate peaks and the peaks of the deconvolved signal. Finally, response traces were smoothed with a Gaussian filter (σ = 31ms).

Electrophysiological signals were high-pass filtered and spike sorting was performed using the CortexLab suite (https://github.com/cortex-lab, UCL, London, England). Single unit clusters were identified using kilosort 2.5 followed by manual corrections based on the interspike-interval histogram and the inspection of the spike waveform using Phy (https://github.com/cortex-lab/phy).

Both for imaging and electrophysiology data, single trial sound responses were extracted (0.5s before and 1s after sound onset) and the average activity over the prestimulus period (0.5s - 0s before sound onset) was subtracted for each trial.

### Reproducibility index and cell selection

To quantify the noise levels in the data, we calculated the mean inter-trial correlation across all pairs of trials. The single neuron reproducibility is then defined for each neuron as the average of the inter-trial correlation for that neuron’s response to all 140 sounds. The population response reproducibility for each sound is defined as the average of the inter-trial correlations of the full sequence of response of the whole neural population to that sound. Region of interests (ROIs) or single units with reproducibility below 0.12 were classified as non-responsive and were excluded from all analyses except population sparseness. As detailed in the **Extended Data Table 1**, the number of responsive units and the corresponding fraction of the total number of units/ROIs recorded are: AC, 19414 (32%), TH, 3969 (12%), THE, 484 (97%), 5936 (39%), 442 (78%).

### Noise-corrected correlation

For each dataset, population representations were estimated after pooling all recording sessions in a virtual population. We used the correlation between population vectors as a metric of similarity between representations. The areas and techniques used to estimate neuronal ensemble representations yielded different levels of trial-to-trial variability due to intrinsic neuronal response variability and measurement noise. Most representation metrics are biassed by variability, even after trial averaging, due to variability residues. For example, the correlation between two population representations (population vectors) will tend to decrease with respect to a variability-free estimate ^43^. When multiple observations of the same representations are available, it is possible to account for the impact of variability, by using specific estimators ^43^. Here we showed analytically (see **Supplementary Information**) that the value of the Pearson correlation coefficient 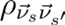 between population vectors for two sounds 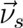 and 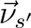 in absence of variability can be exactly estimated from noise-corrupted single-trial observations 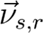 and 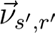 of 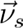 and 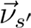 when their dimension N approaches infinity, based on the formula:

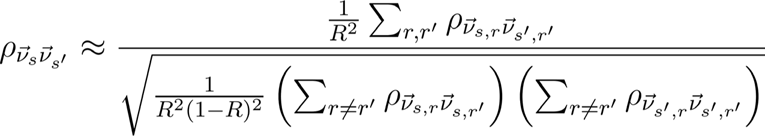

in which r and r’ are single trial indices and R is the total number of trials. This analytical result is confirmed by simulations for finite N, indicating that our estimator converges to the correlation value of the noise-free vectors (**Extended Data Fig. 5c**). Code for calculating this estimator is provided with the online data set.

Simulations for finite N show as expected that the estimator displays substantial deviations around the true correlation which however average to zero. This leads to values of the estimator that can be outside [-1,1] in some cases. Our estimator displays extremely large deviations when 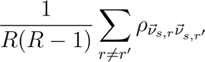 approaches 0, i.e. for representations that are dominated by noise. This occurred more often in datasets obtained by imaging, in particular in the thalamic axonal boutons dataset (TH). To limit imprecisions from these extreme values we excluded from all datasets sounds for which 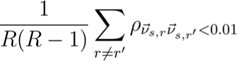. In typical neural data, there are significant noise correlations across simultaneously recorded neurons within a trial. Therefore, the effective N can be much lower than the number of neurons. We minimised this contribution by shuffling trial identity for each neuron independently.

To evaluate the significance of mean correlation differences across all sound pairs for temporal and rate representations, we used a bootstrap procedure over the independently recorded sessions. This procedure had the advantage of providing a statistical assessment for biological replicability based on strictly independent measurements (neurons of the same recording are not fully independent statistically). The noise-corrected correlation measure was estimated 100 times after a random resampling of sessions with replacement. Based on this distribution, we measured the standard deviation and calculated p-values down to 0.01. Sequence correlation was measured on vectors formed by concatenating the responses of all neurons throughout time (vector dimension = N_Neurons_ x N_TimeBins_). Rate correlation was measured first by time-averaging the responses of each neuron and then concatenating these values for all neurons (vector dimension = N_Neurons_). In both cases, we used data from the sound onset to 250ms after the sound offset. To normalise the difference between temporal and rate correlation when comparing between areas we use the formula :

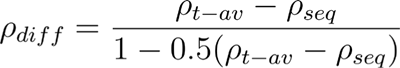

### Noise-corrected sparseness measure

There exist several sparseness measures which are all biassed by variability in neuronal activity measurements ^45,68–70^. The most classical measure as defined in ^68^ is not appropriate for baseline-corrected, linearly deconvolved calcium data because it requires positive response values. We show in the **Supplementary Information** that kurtosis, the 4th order moment of a distribution, is a sparseness measure which can be corrected for variability-related biases and is appropriate for all our datasets. This metric quantifies the “long-tailedness” of the distribution. Sparse response properties correspond to rare and strong responses which generate long-tailed response distributions as opposed to dense response properties which correspond to more compact response distributions. For lifetime sparseness, measured for each neuron separately, Kurtosis is defined as:

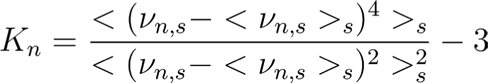

in which <>_s_ indicates averaging over sounds and 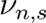 is the noise-free response of neuron *n* to sound *s*. In the case of population sparseness, which is measured for each sound separately, <>_s_ should be replaced by <>_n_ which indicates averaging over neurons. The Kurtosis formula can be developed into the moments of order 1 to 4 of 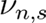.

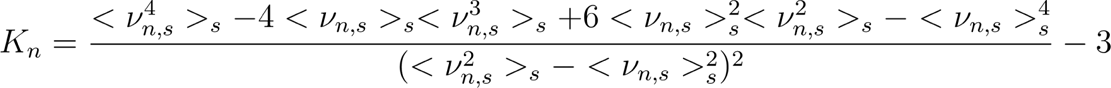

Starting from the second order, estimates of these moments based on trial-averaged response include noise-related bias terms, which skew the kurtosis estimates for limited trial counts. We analytically demonstrated and numerically verified that these biases can be suppressed using noise corrected formulae of all moments that are detailed in the **Supplementary Information**. Code for these calculations is provided with the online data set.

When calculating population sparseness, we analysed all neurons including non-responsive neurons. Non-responsive neurons with aberrant response levels (>5 times the maximal value of responsive neurons) were excluded. Based on this, the percentages of units used were : ICE : 92%, IC: 80%, TH: 61%, THE: 97%, AC:92%).

### Population activity classifiers

To evaluate the accuracy of sound identification based on single-trial population responses, we trained a nearest-neighbour classifier on a subset of trials and cross-validated it on a distinct subset of trials. Training and testing sets were constructed by randomly selecting half of the trials for each unit. For each sound, we correlated the population response averaged over the training trials for this sound with the population response averaged over the testing trials for all the other sounds. The sound with the highest correlation was assigned as the prediction. Decoding accuracy is defined as the proportion of correctly assigned sounds.

Spatial and spatio-temporal were defined as for the correlation measures. Statistical significance was evaluated using the same bootstrap procedure as for the correlation measures. Importantly, decoding depends inherently on trial-to-trial noise which limits the possibility of comparing between areas. This analysis serves to contrast spatial and spatio-temporal codes within an area.

To measure the information contained at different timescales, the temporal sequence of population activity was decomposed into its Fourier coefficients corresponding to a discrete set of timescales ranging from *T*, the 750 ms sound response duration, down to *2Δt,* where *Δt* is the discretization time of the dataset (1/2*Δt = f* the Nyquist frequency; Δt = *T/*24 *=* 31.25 ms for 2P-imaging data and Δt = *T/*96 *=* 7.81 ms for electrophysiology data).

The Fourier coefficient *C_n,r_* for frequency *n/T* and neuron *r* is defined as

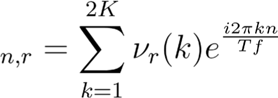

where *ν_r_(k)* is the activity of neuron *r* at timestep *k*, *i* = √−*1* and *K* = *Tf*. Each coefficient is a complex number or, equivalently, a two-dimensional vector. Hence the activity sequence for a given neuron is either represented by a vector of *2K* data points or of *2K* Fourier coefficients.

To measure the information present at a given time scale, we applied the population activity classifier on the population vector containing the 2N Fourier coefficients for this time scale for the N neurons of the dataset (**Extended Data Fig. 5e**). To measure information present above a particular time scale T_max_, we used the Fourier coefficients from 1 to T_max_ for each neuron and concatenated them into a 2NT_max_ population vector (**Fig. 2k**). Of note, when evaluating information at particular time scales, we did not apply any temporal filtering steps to avoid artefacts due to the finite size of the filter and preserve the full bandwidth of the data.

### Tuning analysis

To quantify the number of neurons significantly tuned to a specific property, we first performed a parametric ANOVA test to identify the neurons which respond significantly more to one of the sounds of interest (e.g. 60, 70 or 80 dB levels across all pure tones for intensity tuning, up vs down modulations in a given frequency range for frequency modulation). We used a threshold of p=0.05. We do not compare the absolute number of neurons tuned to a given property between areas since this will largely reflect the different levels of noise in the data sets and we focus on the properties of significantly tuned neurons.

To measure the tuning of individual units to classes of stimuli (for example up chirps vs down chirps) we used the following modulation index:

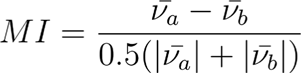

### Reinforcement learning model

We adjusted a previously published reinforcement learning model ^27,49^, to learn discriminations between pairs of temporal inputs. The model receives as inputs the temporal responses for two sounds: (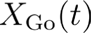) for the rewarded sound and (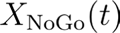) for the non-rewarded sound. The model learns the synaptic weights between these input representations and a downstream decision circuit (**Fig. 3a**). This circuit is composed of a Go-unit which outputs the decision (synaptic weights :) and an inhibitory neuron that provides immediate linear inhibition to the reward neuron (synaptic weights : *W_I_*). The temporal output, *y(t)*, of the model can therefore be described as :

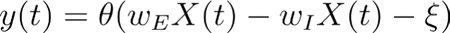 where *θ* is the Heaviside step function, ξ is a time - independent Gaussian random noise process that models stochasticity of behavioural choices. The decision to go is made if the mean activity of the Go-unit within the response window 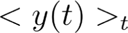 is larger than 0.2 (< · >*_t_* denotes time averaging over 0.5s).

The synaptic weights are updated according to a learning rule which compares the reward prediction to the actual reward, assuming that reward prediction corresponds to the mean input received by the Go-unit. The learning rule has three particularities that have been previously shown to be important to account for mouse behaviour ^49^ and compatible with our knowledge of synaptic plasticity rules. First, it is asymmetric : the learning rate is larger when an unexpected reward occurs than when an expected reward does not. Second, it is multiplicative: the learning rate at a given synapse depends on the current weight of that synapse. Finally, it takes into account the known dynamics of the eligibility trace in the striatum ^52^ which is a key target of both AC and TH in discrimination learning ^33^. The eligibility trace is a key mechanism in the “neo-hebbian framework” that aims to explain how synaptic plasticity can accommodate delays between action initiation and environmental feedback. This theory proposes that synapses that undergo pre-post coincidence prior to feedback are tagged via a long-lasting (∼ few seconds) eligibility trace. Weight changes will only occur at these tagged synapses if they are subsequently exposed to neuromodulatory feedback before this eligibility trace decays. In line with this, in the striatum, potentiation of synapses is conditioned on dopamine release within a ∼3s time window following coincidence of pre- and post-synaptic activity ^52^. To implement this in our model, the temporal signal for the model input is convolved with a kernel corresponding to the temporal profile of dopaminergic plasticity gating taken from Yagishita et al ^52^ before calculation of the weight update.

The learning rule is implemented as :

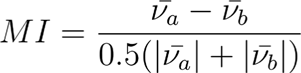

Where λ the learning rate, *R* is the action outcome (*R* = 1 for reward, *R* = −1 for no reward, σ is the behavioural noise level parameter that sets the models peak performance, *f*() is the function that implements asymmetric learning such that

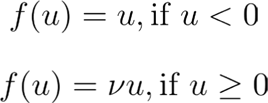

*v* > 1 is the learning rate asymmetry ratio,

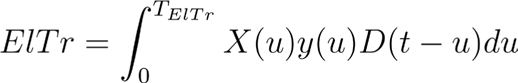

where *D*(*u*) is the temporal function shown in **Fig. 3a** and taken from Yagishita et al ^52^ and *T_ElTr_* = 0.5s.

In order to estimate the speed at which the model learns to discriminate between different neural representations, we used as input the population vector time series for two different sounds from a given area. For calcium imaging, we first performed clustering of the response to reduce dimensionality. The model was then run for three independent simulations to average out the stochastic contribution and we evaluated the number of trials to reach 80% based on the average learning curve over these three repeats.

For dimensionality reduction of the population vector, we performed agglomerative hierarchical clustering based on the euclidean distance between each neuron’s full temporal response to all stimuli. The number of clusters was established by increasing the number of clusters until the sound-pair RSA matrix constructed from the clusters explained 95% of the variance of the matrix constructed from the full neural population. Clustering was performed independently for each data set and yielded approximately 150 clusters in all areas. AC data displayed in **Fig. 2c** represents clusters rather than single neurons.

### Convolutional neural networks

#### Augmented sound set

In order to train deep neural networks, we created an augmented sound set that covered all the basic parameters explored by the original 140 sound set used in experiments. We first augmented the basic sounds composing the sound set from 140 to 2169. This first step generated the sounds by independently varying all features defining the sounds (frequency, intensity, amplitude modulation direction or period, frequency modulation direction, chord composition). Thereby, a given feature cannot be predicted based on other features as in the experimental sound set. We further augmented the sound set using the approach from ^55^. Each 500ms sound is embedded at a random time in a randomly chosen 1.5 s snippet taken from an auditory scene (bus station, park, street…) with a random intensity (average : 53db, std : 7dB). We thus generated a total of 150.000 sounds for the test (6.000), train (110.000) and validation (34.000) sets respectively.

#### Task definitions

The multi-category task required the network to output a 14-element binary category vector in which 1 indicates that the sound presented belongs to one of 14 categories, divided into 4 groups within which categories are mutually exclusive: frequency range, intensity range, frequency modulation type, and amplitude modulation type. However, all sounds had to receive one label from each group. The group structure was not provided to the network which therefore had to learn that a sound could not be simultaneously high and mid frequency for example. The categories were defined as follows:

- Frequency range group: high frequency (4-8 kHz) / mid frequency (9-17 kHz) / low frequency 18-38 kHz) / broadband (white noise only). For chords and frequency modulated chirps, the frequency value used for categorization was the average of all frequencies (i.e. middle of the chirp).
- Intensity range group: high time-averaged intensity (80dB) / mid time-averaged intensity (70dB) / and low time averaged intensity (60 dB). Amplitude modulated sounds were assigned to their closest time-averaged range group. We obtained different overall intensities by ramping sounds sublinearly, linearly or supralinearly.
- Amplitude-modulation group: Up-ramping/ down-ramping / sinusoidal modulation / no modulation.
- Frequency-modulation group: Up chirp / Down chirp / no modulation.

The sound identification task required the network to output the identity of each of the 2169 different sounds without any category.

The convolutional autoencoder is a network trained to reproduce with minimal loss its input with the constraint of passing all information through a small, central bottleneck layer. It is composed of an encoder sub-network that processes the input to allow for compression in the bottleneck layer and a decoder sub-network that reconstructs the output from the low-dimensional bottleneck representation.

#### Architecture definition and training

All networks take as input a 2D (time x frequency) matrix of the log-scaled spectrogram of the sound and must produce as output the labels described above. In order to achieve this, a series of convolutional blocks is applied to transform the input. All classification networks were built from a series of 6 blocks composed of the same layers :

- convolution : the input is convolved by a filter whose weights the network must learn, each layer applies multiple filters, generating a 3D matrix (time x frequency channel) from the initial 2D input (free parameters : kernel size, kernel stride, channel number)
- activation : the output of the convolution is passed through a Relu non-linear activation function
- maxpooling : the output of activation is downsampled by taking the maximal value of neighbouring values (free parameters : pool size, pool stride)
- dropout : in order to improve the robustness of training, during each training batch a random 50% selection of connections are eliminated. During testing and validation, all connections are active.

After these convolutional blocks, a final 64-node fully connected layer with a Relu non-linearity allows to aggregate information across time, frequency and channel dimensions. The output layer is obtained for the multilabel task by applying a sigmoid function to the fully connected output and for the identification task by applying a softmax function.

The output of the last layer allowed us to calculate the value of the loss function that comprises the error the network makes (categorical cross entropy loss function) and a L1 regularisation term in order to improve network robustness. This loss was then back-propagated during training in order to optimise the weights of the connections using the Adam optimizer.

Any given architecture requires arbitration across a wide range of free parameters, most notably the kernel and max pooling size and stride as well as the number of channels in each block. One approach to this problem is to perform a search across architectures to obtain optimal performance on the task. This has allowed optimization on ecologically-relevant tasks to be proposed as a criteria for building deep networks that function like the brain. However we focused on general properties of CNNs and were using a simple task without natural sounds. We therefore chose to assess the generality of our results on various architectures instead of performing an exhaustive search. We also verified the reliability of our results for a given architecture by using 2 different initialization weights per architecture. The four architectures we evaluated are defined as follows (CV : convolution layer, MP : max pooling layer, FC : fully connected layer, Ker : kernel size) :

1. Input : 109 x 150; Cv1 : 109 x 150 x 18, Ker(3,3); MP; CV2 : 55 x 75 x 20, Ker(5,5); CV3 : 55 x 75 x 24, Ker(6,6); MP; CV4 : 28 x 38x 28, Ker(7,7); CV5 : 28 x 38 x 32, Ker(8,8); MP; CV6 : 14 x 19 x 32, Ker(9,9); FC : 64
2. Input : 109 x 150; Cv1 : 55 x 75 x 18, Ker(3,3); CV2 : 55 x 75 x 20, Ker(5,5); CV3 : 28 x 38 x 24, Ker(6,6); CV4 : 28 x 38x 28, Ker(7,7); CV5 : 14 x 19 x 32, Ker(8,8); CV6 : 14 x 19 x 32, Ker(9,9); FC : 64
3. Input : 109 x 150; Cv1 : 55 x 75 x 1, Ker(7,7)8; CV2 : 55 x 75 x 20, Ker(7,7); CV3 : 28 x 38 x 24, Ker(7,7); CV4 : 28 x 38x 28, Ker(7,7); CV5 : 14 x 19 x 32, Ker(7,7); CV6 : 14 x 19 x 32, Ker(7,7); FC : 64
4. Input : 109 x 150; Cv1 : 55 x 75 x 24, Ker(3,3); CV2 : 55 x 75 x 24, Ker(5,5); CV3 : 28 x 38 x 24, Ker(6,6); CV4 : 28 x 38x 24, Ker(7,7); CV5 : 14 x 19 x 24, Ker(8,8); CV6 : 14 x 19 x 24, Ker(9,9); FC : 64

One prominent consequence of the choice of CNN architecture is the way in which the input volume evolves throughout the network. Choosing a large stride in the convolutional or a large window size in the max pooling layer will lead to a shrinkage of the input dimensions (time and frequency). Given that the temporal dimension is preserved in the brain, we examined an architecture in which there is no shrinkage at all of the temporal dimension. To do this, we used the 4 same architectures described above, with the temporal dimension kept constant by setting all strides to 1 and eliminating max pooling. This results in a large expansion of the parameters in the network and affects training speed although asymptotic performance remains the same (**Fig. 4b**).

The convolutional autoencoder receives as input the 2D spectrogram and must output a denoised spectrogram (spectrogram of the central sound without the background noise). The autoencoder was composed of 4 convolutional blocks as previously described in the encoding part and decoding networks, the bottleneck is a fully-connected, 20 node layer. Training was performed with an Adam optimizer, L1 and L2 regularisation and MSE as a loss function.

The convolutional neural network trained on word and musical genre recognition was previously published^55^ and parameters have been made available at (https://github.com/mcdermottLab/kelletal2018). This network is composed of a central branch that splits into two branches, with one branch trained to identify musical genres and the other branch trained to identify words. In the original paper, the network was shown to achieve human-like performance and to qualitatively reproduce psychophysical measures during these tasks.

#### Analysis of CNN activations

Once the networks had been trained, we analysed the responses of all nodes in each activation layer to the 140 sounds that were presented during experimental sessions. Each sound generates at a given layer a 3D matrix (time x frequency x channels). By considering the temporal response of each frequency x channel combination we obtained analogs to the temporal response of individual neurons. We then applied the same analysis techniques to these artificial responses as described above for neural recordings. In order to perform decoding which requires multiple presentations of the same sound, we presented to the network multiple copies of each sound embedded in different noise backgrounds.

### Cochlear model

A computational model was implemented by adapting the seminal model of Meddis ^71,72^ to the mouse cochlea and validating it with mouse auditory nerve recordings ^73^. The model consists of a cascade of six stages recapitulating stapes velocity, basilar membrane velocity, inner hair cell (IHC) receptor potential, IHC presynaptic calcium currents, transmitter release events at the ribbon synapse, and firing response in auditory nerve fibres (ANFs) including refractory effects. The input model is a sound stimulus (in Pascals). The output is a train of spiking events (in spikes/s) in 590 ANFs innervating 40 IHCs with a characteristic frequency (CF) distributed at regular intervals along the cochlear tonotopic from 5 to 50 kHz, 12 IHCs per octave. This distribution covered 82.8% of the basilar membrane length from 1.2% (apex) to 83.9% (base) in 2.07% increments. According to experimental data, the number of ANFs per IHC (N) was controlled by the relationship N=−0.0038x^2+0.375×+7.9 where x is the IHC location along the basilar membrane such that x=−56.5+82.5 log⁡(CF), with x in percent from the apex and CF in kHz. By adjusting the time constant of the calcium clearance τ_Ca within each IHC synapse, ANFs with different spontaneous discharge rate (SR=91.1 *τ*_Ca_^2.66^, with *τ*_Ca_ in ms and SR in spikes/s) were simulated from 0.5 to 95 spikes/s (21 ± 19.8 spikes/s, mean ± SD) to match the SR distribution reported in mouse auditory nerve.

### Fast time-scale temporal to rate conversion in an excitation / inhibition model

We modeled the response of two integrate-and-fire neurons connected by a single inhibitory synapse that receive inputs A (In_A_) and B (In_B_) respectively. These inputs represent the A and B driven populations in the optogenetics experiment of **Fig 1**. The A and B inputs exactly reproduced our temporal-coded stimulation (225ms duration, 20Hz stimulation, 25ms between flashes) but we systematically varied the interval between A and B which in the experiment was set to 250ms. The excitatory neuron received excitatory input A input of synaptic strength (J_A_=0.09) and inhibitory input from the inhibitory neuron (In_I_) of synaptic strength (J_I_=0.04), delayed relative to inhibitory spiking (*τ*_I_=2ms). The inhibitory neuron received excitatory input B of synaptic strength (J_B_=0.09). Both neurons decayed to their resting membrane voltage (V_m_=−65mV) with membrane constant (*τ*_M_=10ms), consistent with *in vivo* findings ^38^ and contained a white noise term (In_noise_). They emitted a single spike when they reached the threshold voltage (V_T_=−50mV) and their voltage was then reset to (V_R_=−70mV).

The equation of the voltage for each neuron is given by :

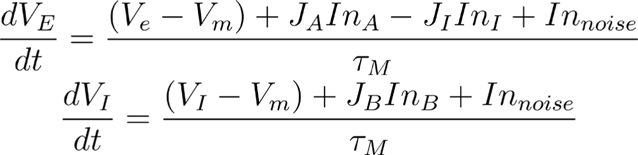

### Statistical analysis

Statistical results (degrees of freedom, p-values and statistical values) are reported in figure legends or in **Extended Data Table 3**. For statistical analysis of neural data, we performed a bootstrap analysis as detailed above. For statistical analysis of behavioural data provided in the manuscript, the Kolmogorov–Smirnov normality test was first performed on the data. If the data failed to meet the normality criterion, statistics relied on non-parametric tests. We therefore represent the median and quartiles of data in boxplots in all figures, in accordance with the use of non-parametric tests. Ranksum and signed rank: we report the signed rank statistic if the number of replicates is too weak to provide the normal Z statistic.

#### Acknowledgments

We thank Maia Brunstein of the Hearing Institute Bioimaging Core Facility of C2RT/C2RA for help in acquiring Airyscan images of thalamocortical boutons in the auditory cortex and Alexander Kell for help implementing the word and music classification network. We also thank Yves Boubenec, Yves Frégnac, Andrew King, Srdjan Ostojic and Christine Petit for their feedback on the manuscript. We acknowledge the support of the Fondation pour l’Audition to the Institut de l’Audition.

We acknowledge the support of the following funding sources: Fondation pour l’Audition, FPA IDA02 (BB) and APA 2016-03 (BB) European Research Council, ERC CoG 770841 DEEPEN, (BB) Fondation pour la Recherche Médicale SPF202005011970 (SB) European Union’s Horizon 2020 research and innovation programme under grant agreement No 964568, project Hearlight (BB)

## Author contributions

S.B., Ja.B., A.K. and B.B. conceived experiments, designed the study and interpreted data. S.B., Ja.B., A.K., T.T., J.S., A.V. and B.B. collected data and S.B., Ja.B., A.K., S.C. and B.B. performed data analysis. Je.B. and J.L.P conceived and implemented the cochlear model. B.B. and S.B. implemented the reinforcement learning model. S.B., K.B. and Y.G. implemented the deep learning models. S.B., Ja.B. and B.B. prepared figures. S.B. and B.B. wrote the manuscript. S.B and B.B. managed the project.

## Materials & Correspondence

All datasets are freely available at 10.12751/g-node.sz67di, hosted by G-Node Infrastructure. Custom codes used in this study are freely available at 10.12751/g-node.sz67di, hosted by G-Node Infrastructure. Further requests should be addressed to

## Supplementary Information

### Supplementary information about the auditory system dataset

To rapidly obtain large datasets from these structures, we used GCAMP6s-based two-photon calcium imaging of either cell bodies (AC and IC, **Extended Data Fig. 3a & d**) or axonal projections (TH, imaged in AC) (**Extended Data Fig. 3b**). Collecting data simultaneously from around 1000 AC neurons or TH axonal boutons and from 100 to 200 neurons in IC, we could extensively sample representations in each region. In AC, all 60.822 ROIs were mapped to functional subfields based on tonotopic gradients ^74^ and to the cortical layer from imaging depth (**Extended Data Fig. 4a-f).** 70% of ROIs were in primary auditory cortex (A1), the largest subfield of AC, but the anterior, suprarhinal and dorsal posterior auditory fields were also covered (**Extended Data Fig. 3a & 4e**). Moreover, with recording depth reaching up to 600 µm, we sampled neurons from layers 1 to 5 with an emphasis on layers 2 and 3 (**Extended Data Fig. 4f**). Therefore, with the exception of layer 6 and of the small ventro-posterior subfield, the whole of primary and secondary AC was extensively covered. Inputs from TH were sampled with 39.191 putative TH axonal boutons spread across AC (75% of ROIs in A1) (**Extended Data Fig. 3b**) and validated post-hoc with the thalamic marker VGLUt2 (**Extended Data Fig. 4g,h**)^75^. In addition, we recorded 15.132 ROIs in the dorsal IC down to 250µm depth (**Extended Data Fig. 3d**).

Since calcium imaging and deconvolution has not been verified for TH axons, we performed electrophysiological recording in primary and secondary auditory TH (498 single units, **Extended Data Fig. 3c**). Electrophysiology was also used to cover the central inferior colliculus (563 single units), the main primary subregion of this structure (**Extended Data Fig. 3e**). Electrode locations were identified with post-hoc histology and short-latency responses (**Extended Data Fig. 3c,e**). Finally, we used a detailed biophysical model of the cochlea calibrated against auditory nerve recordings ^73^ (AN), to provide insight into the information entering the auditory system (**Extended Data Fig. 3f, 4i,j**).

Calcium signals were temporally deconvolved using a linear algorithm to retrieve estimates of neuronal firing rate variations that are robust to parametrization errors ^76^. This allowed us to reach a ∼150 ms temporal precision as estimated from responses to amplitude modulated sounds (**Extended Data Fig. 3a,b,d**). The temporal modulations of our sounds were chosen to evolve at timescales compatible with this resolution of calcium imaging. This was confirmed by our decomposition of neural population activity into specific timescales using Fourier analysis (**Extended Data Fig. 5e**). This revealed that even with electrophysiology, in which activity contained information at fast timescales up to 30Hz, information nonetheless saturated at around 3Hz. Therefore all information needed to discriminate our sounds is available below 3Hz, which matches calcium imaging resolution, with information at faster timescales being redundant.

## Extended Data Figures and Tables

**Extended Data Fig. 1.**
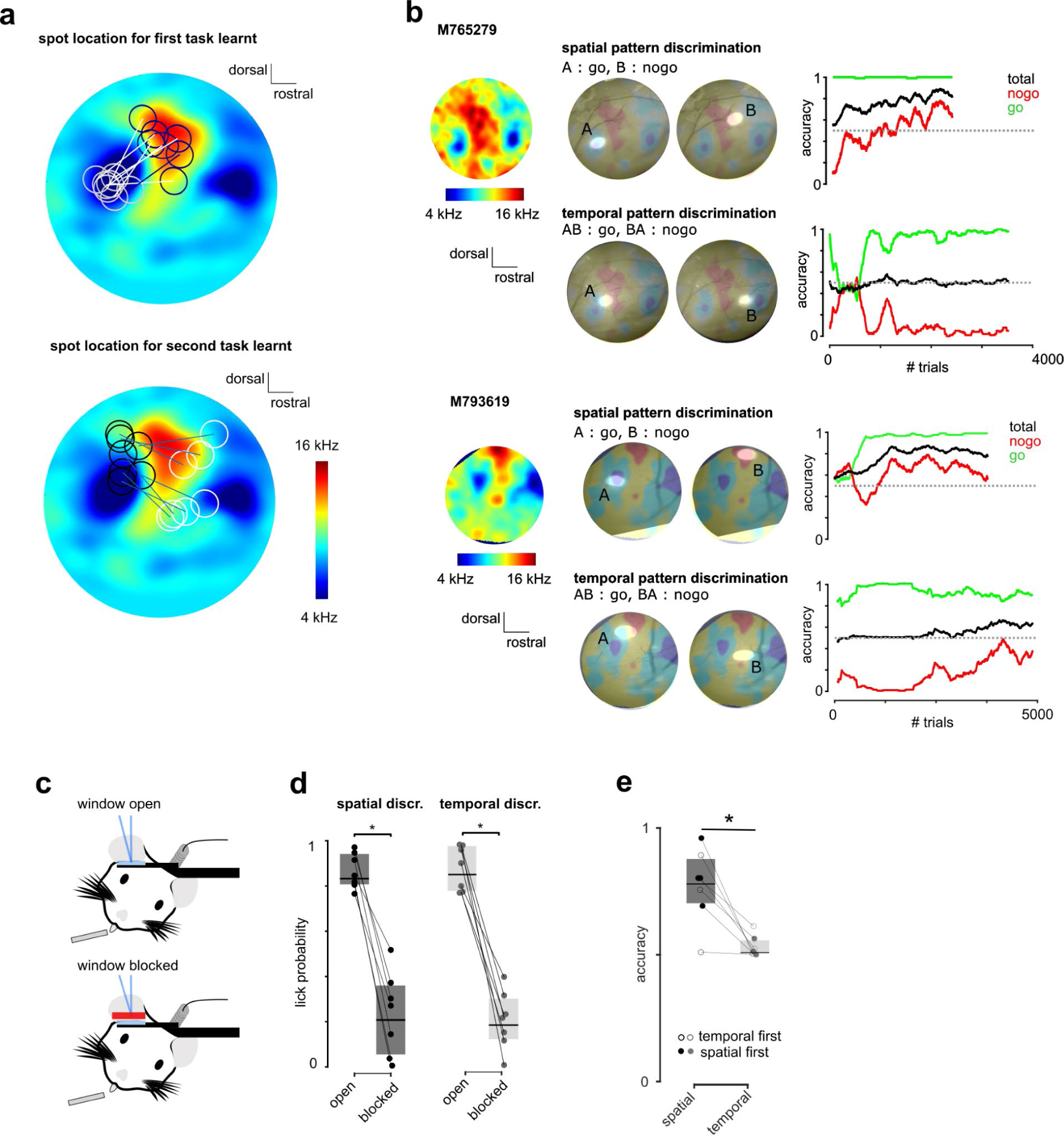
Details of behavioural learning in optogenetic cortical stimulation protocol. **a.** Population average intrinsic imaging map of tonotopic areas in AC showing the localization of all spots used for optogenetic stimulation (n=7 mice). **b.** Intrinsic maps and spots used for stimulation with learning curves from two example mice in both tasks. **c-d.** Control experiment showing that response to optogenetic stimulation is specific to cortical activation : mice ceased responding to light stimulation when the cranial was blocked by a small cache that left all other light cues intact. Note also that the lick probability for temporal and rate patterns is identical during this initial phase. (paired Wilcoxon test, p = 0.0156, signed rank value = 28, n=7) **e.** Accuracy over the last 300 trials for all mice. (paired Wilcoxon test, p = 0.015, signed rank value = 28, n=7).

**Extended Data Fig. 2.**
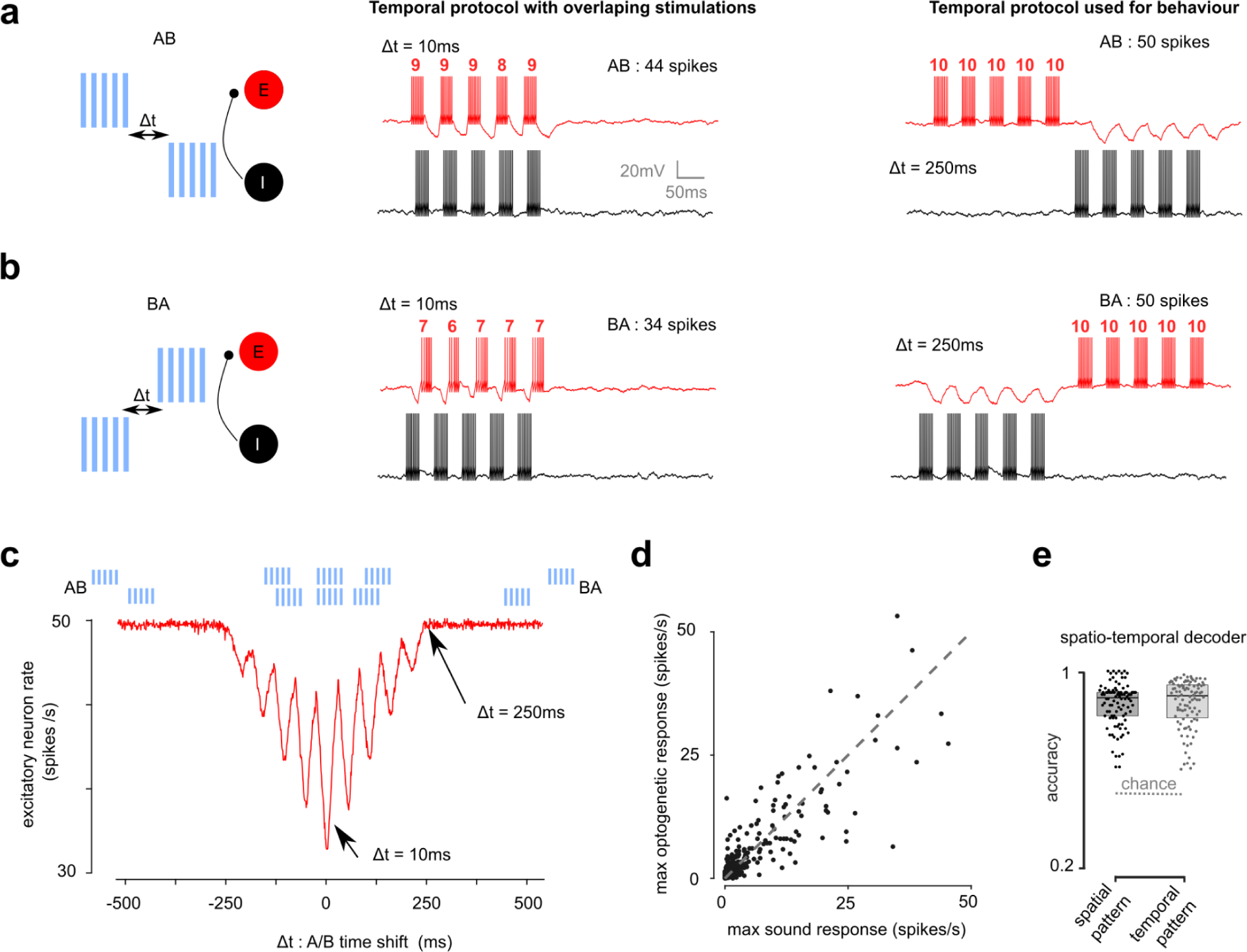
Synaptic integration converts temporal information into firing rates at short time scales. Neurons integrate synaptic inputs over the timescale of their membrane time constant. Hence the number of output spikes depends not only on the number and size of synaptic inputs but also on their relative timing. This property is reinforced by synaptic connectivity such that simple circuits can detect time shifts between incoming inputs and thereby transform temporal information into firing rate information. **a.b** To illustrate this point and evaluate in which conditions temporal information injected in the AC may give rise to salient firing rate representations, we simulated the activity of an integrate-and-fire neuron E connected to a neuron I by an inhibitory synapse. The inputs received by E and I are trains of 5 current pulses at 20Hz (pulse duration 25ms, total duration 225ms) simulating Chr2 activations by light as in Fig. 1. Input onsets to neurons E and I are shifted by a time Δt to generate temporal information. Exemple simulations indicate that when the time shift is small and the inputs to E and I overlap in time (middle panels), neuron E emits fewer action potentials than when the inputs are well separated in time (right panels). **c.** Quantification of the effect exemplified in panels **a** and **b**, showing the firing rate of neuron E for a range of time shifts Δt. For |Δt| < 250ms, neuron E emits fewer action potentials than for |Δt| > 250ms and the number of action potentials depends on Δt. Hence, temporal information is converted partially to rate information for |Δt| < 250ms but not for |Δt| > 250ms. The boundary value for |Δt| (here ∼250ms) depends on the membrane time constant of the neurons (here set to 10ms, similar to values reported *in vivo* ^38^). In general, this simulation indicates that in order to avoid the conversion of temporal information into rate information, the temporal sequences injected in cortex must avoid temporal contiguities over the time scale of the membrane time constant. Moreover, the use of time-reversed sequences tends to ensure smaller firing rate differences across sequences, compared to time sequences that are not symmetric of each other. **d**. Mean firing rate during the optogenetic stimulation and sound stimulus that evoked the highest firing rate in each neuron (n=321 units). **e.** Accuracy of neural decoder trained to discriminate the patterns used in the task with all spatial and temporal information available in the population vectors. (n=321 units, bootstrap over units, p-value of accuracy vs chance level of 0.5: 0.01, 0.01)

**Extended Data Fig. 3.**
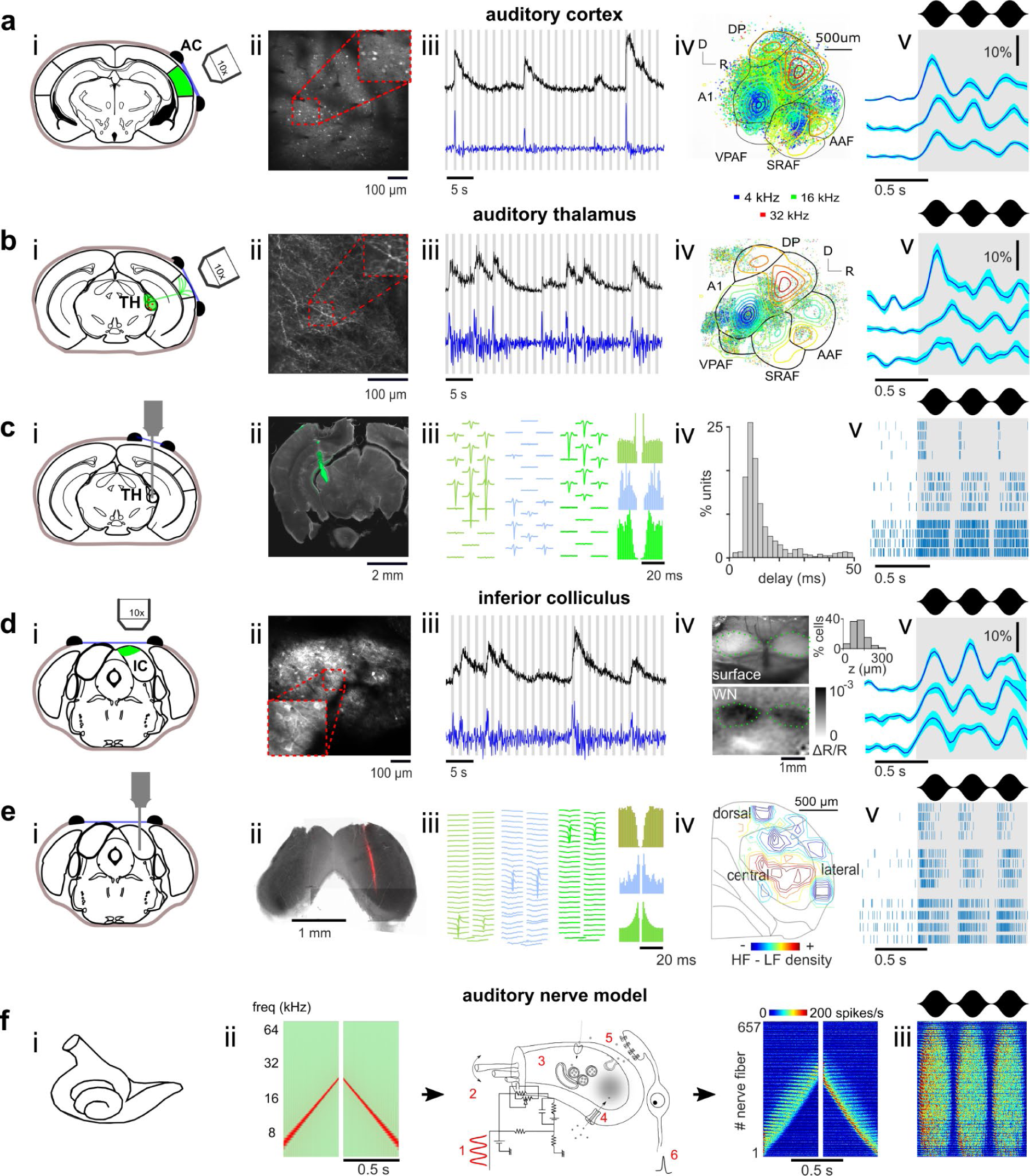
Extensive neural recordings throughout the auditory system. **a.** (i) Schematic of imaging strategy, (ii) sample field of view, and (iii) raw (black) or deconvolved (blue) calcium traces (grey bar: sound presentation) for a sample neuron in AC. (iv) Location of all recorded neurons, colour-coded according to their preferred frequency at 60dB, overlayed with the tonotopic gradients obtained from intrinsic imaging. (v) Response of 3 neurons to 3Hz amplitude modulated white noise. **b.** Same as in **a** for thalamic axon imaging. **c.** (i) Schematic of recording strategy, (ii) sample histology with di-I strained electrode track, (iii) average waveforms and auto-correlograms of three single units, (iv) response latencies of all single units, (v) raster plot of 5 trials from 3 sample units in response to 3Hz modulated white noise. **d.** Same as **a** for dorsal IC except for (iv): view of the cranial window and intrinsic imaging response to white noise. Inset histogram shows distribution recording depths. **e.** Same as **c** for central IC, except for (iv): reconstructed of IC tonotopy from single units. **Ff** (i) Schematic of the cochlea and (ii) of the biophysical model taking a sound as input and providing the responses of auditory nerve fibres. (iii) Response to 3Hz amplitude-modulated white noise. A1 : primary auditory cortex, DP: dorsal posterior field, AAF: anterior auditory field, VPAF : ventral posterior auditory field, SRAF : suprarhinal auditory field.

**Extended Data Fig. 4.**
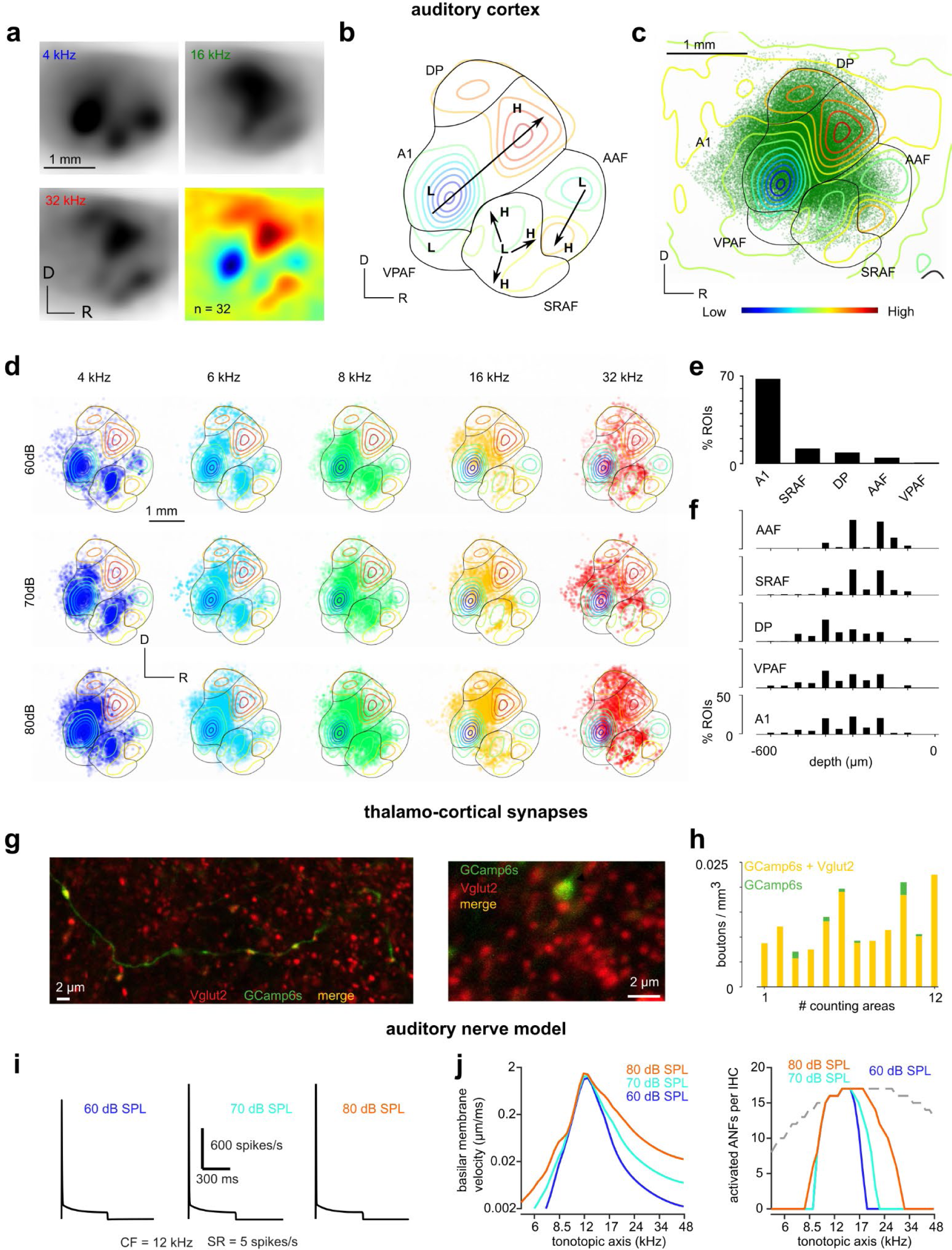
Details of auditory system sampling. **a.** Mean intrinsic imaging responses (n=32 mice) for 4, 16 and 32 kHz sounds (black) and the subtraction of 32kHz and 4kHz maps (colour). This extended data set allowed us to construct a consensus map to align mice included in the study. **b.** Illustration of method used to identify AC subregions based on the tonotopic gradients established in ^74^. **c.** Localization of all recorded ROIs on the consensus tonotopic map with AC subregions. **d.** Localization of responsive neurons to increasing frequency and intensity. Note the larger recruitment with stronger intensity and the spatial shift with frequency. **e.** Proportion of units per subarea. **f.** Depth distribution of units per subarea. **g**. Example thalamocortical axon expressing GCaMP6s merged with Vglut2. Thalamic axonal boutons expressing Vglut2 appear yellow as shown in the magnified region (right). **h.** Density of labelled boutons (Vglut2^+^;GCaMP6s-expressing in yellow; GCaMP6s alone in green) in layer 1 of the AC (12 sample regions; 4 regions per confocal image; means and STD: 0.0122±0.0052, 0.0005±0.0008, density of co-labelled and green only boutons, respectively). **i**. Peristimulus time histogram of an auditory nerve fibre (ANF) with a characteristic frequency equal to that of the presented 12-kHz tone burst (10-ms rise/fall, 500-ms duration) with increasing level from 60, 70 and 80 dB SPL. Note the rapid adaptation of the firing. **j**. Basilar membrane velocity and sound-activated auditory nerve fibres per inner hair cell (IHC) along the tonotopic axis. Note the reduced frequency selectivity with the increasing intensity. Gray dashed line shows the mouse synaptic cochleogram. The criterion for sound-activated auditory nerve fibres was 10 spikes/s above the spontaneous rate.

**Extended Data Fig. 5.**
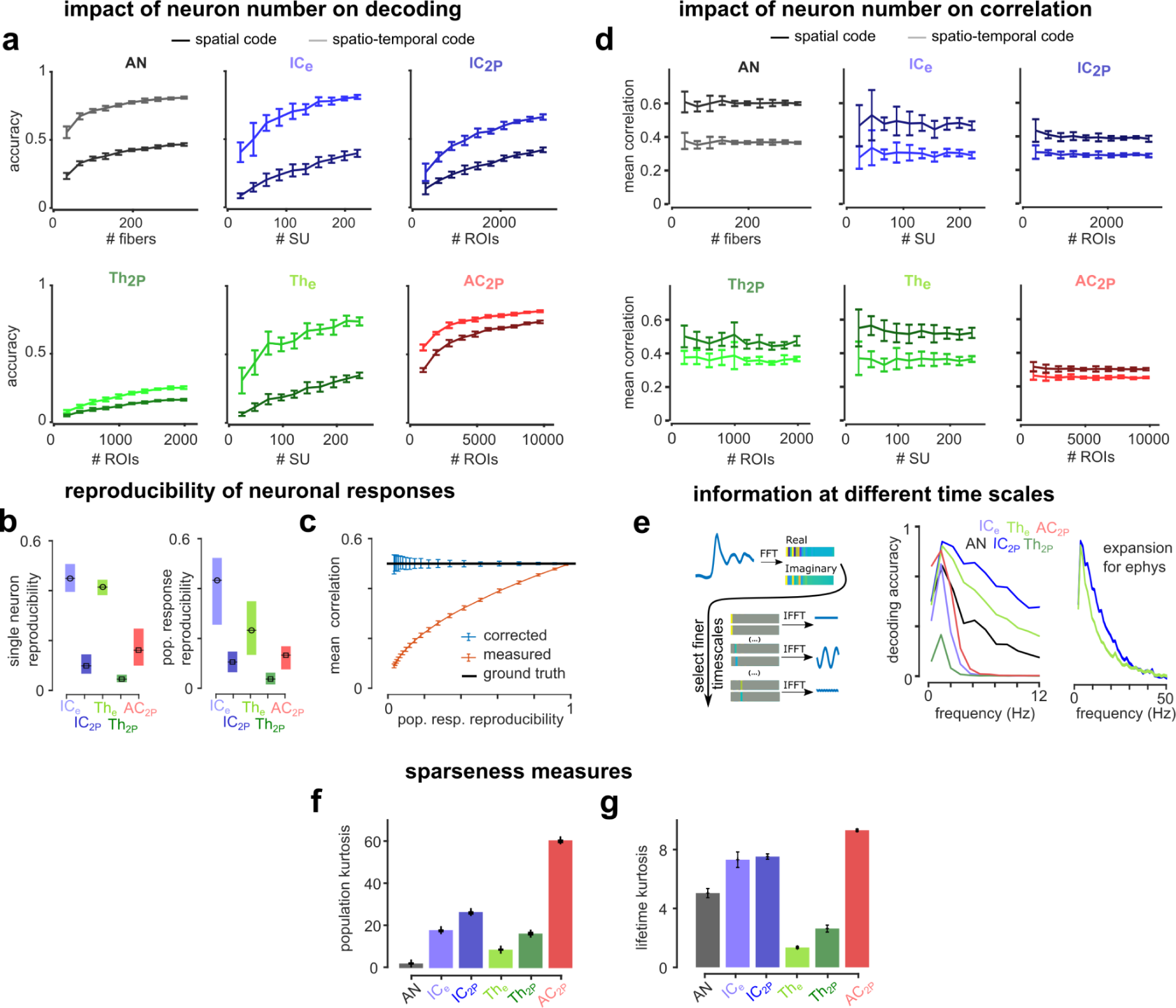
Robustness of accuracy and similarity measures. **a.** Decoding accuracy for spatio-temporal and spatial codes in each area with varying numbers of sub-selected neurons. **b.** Reproducibility of single neuron (left) or population (right) responses measured as the mean inter-trial correlation between responses across sounds (left : n=number of neurons per area, right : n=140 sounds, error bars are quantiles). **c.** Measured correlation of simulated data with low to high response reproducibility before (orange) or after (blue) noise-correction. **d.** Noise-corrected correlation for spatio-temporal and spatial code in each area with varying numbers of sub-selected neurons. **e.** Sketch illustrating the decomposition of population responses by timescale and mean decoding accuracy based on successive Fourier coefficients of neural responses. 0 Hz = spatial code. As expected, 2-photon data only contained information up to whereas electrophysiology data was informative even up to 30 Hz. **f-g.** Noise-corrected sparseness measured using kurtosis. n = 140 sounds for population kurtosis (**f**) and n = ‘all neurons’ for lifetime kurtosis (**g**).

**Extended Data Fig. 6:**
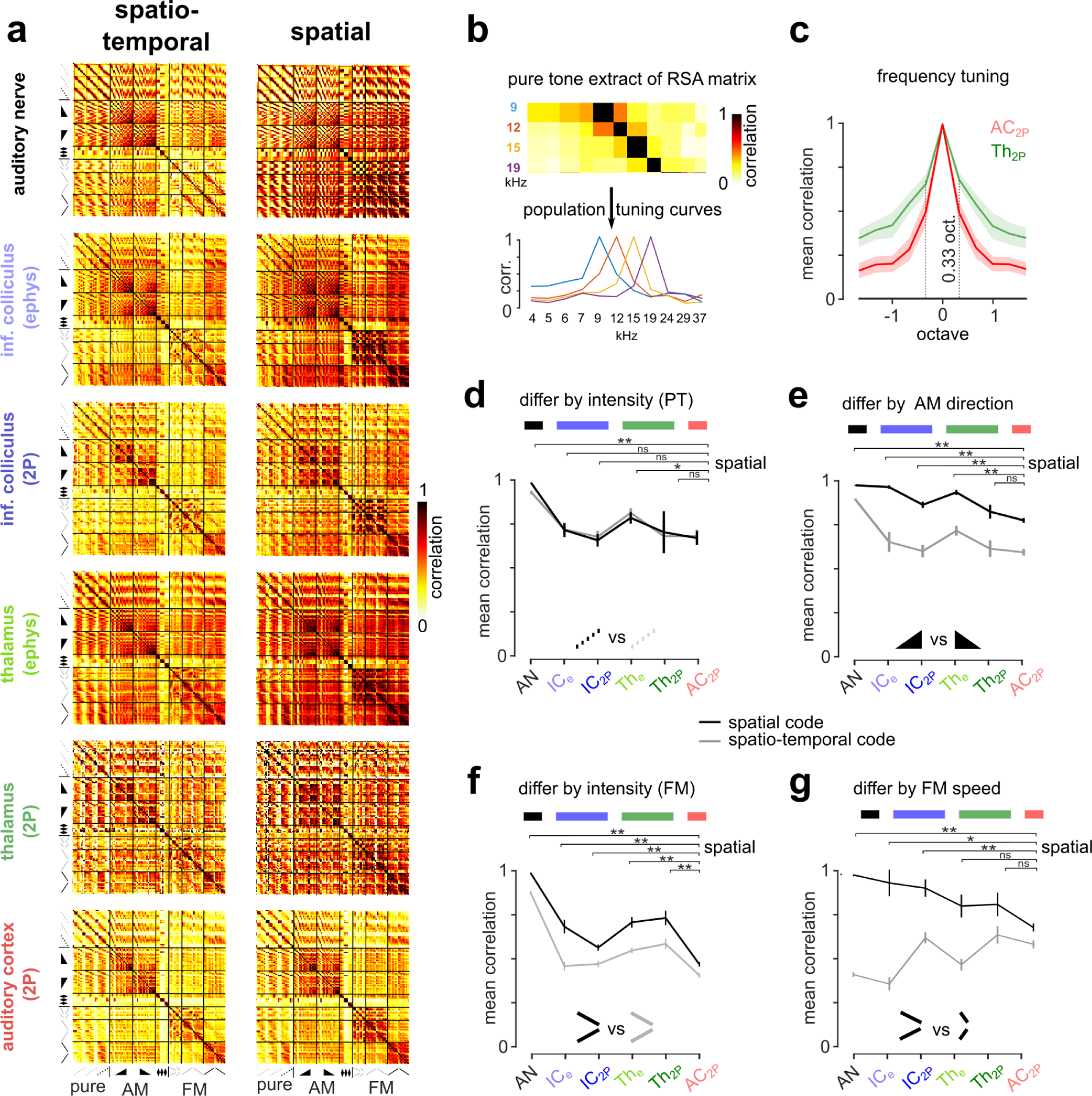
Time-independent rate representations of time-symmetric sounds decorrelate in AC. **a.** Noise-corrected RSA matrices for all sound pairs for temporal (left) or rate (right) codes. **b.** Illustration of method to calculate population tuning curves shown in B from RSA matrix. **c.** Mean noise-corrected correlation between pure tones as a function of their frequency separation. **d–g.** Mean noise-corrected correlation between sound pairs differing by only one acoustic property. **d**. Pure tones at the same frequency differing by intensity, **e**. amplitude ramps at same frequency differing by direction. **f.** frequency sweeps with identical frequency content and duration at 60dB vs 80dB, **g**. frequency sweeps with identical frequency content of different duration, For sounds without temporal structure, correlation of representations are similar in AC and IC, whereas for time-symmetric sounds, all brain areas show larger rate correlations than in the cortex, except for TH2P in **e** likely due to the high variability of thalamic responses. p-value for 100 bootstraps comparing rate correlation of each region to AC, error bars are S.D. Statistical test details are given in the **Extended Data Table 3**.

**Extended Data Fig. 7.**
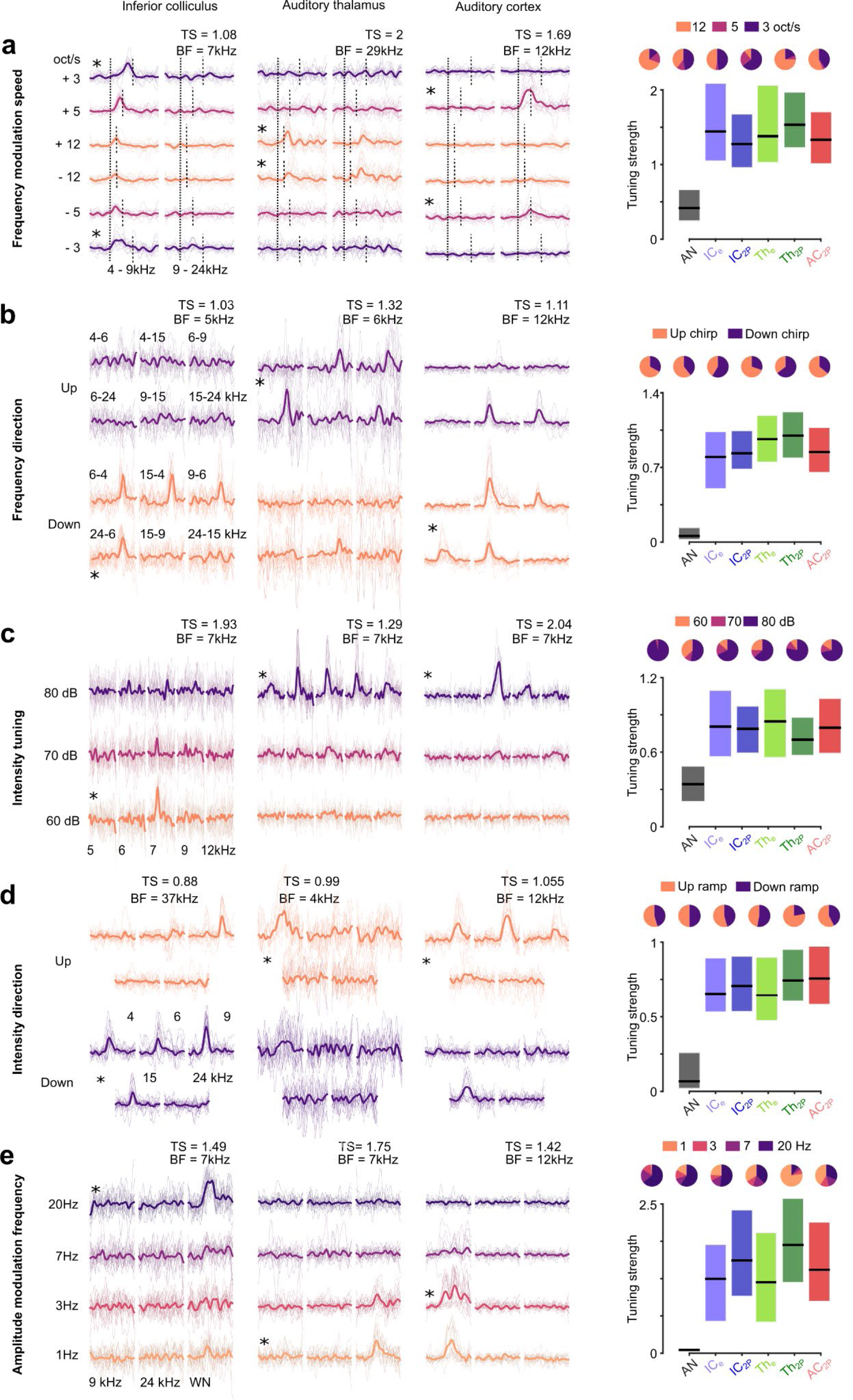
Single cell tuning to diverse acoustic features from cochlea to auditory cortex. **a-e. Right:** For each tuning property we show the responses of example neurons from the IC, TH and AC to sounds that differ according to that property and provide the tuning strength (TS) and best frequency (BF) for that neuron. Asterisks indicate significant tuning of the neuron to a specific value, for example the leftmost neuron in **a** is an IC neuron that is significantly tuned to frequency modulation speed with a maximum response for decreasing frequency at 3oct/s. (left) Boxplot giving the distribution of tuning strengths across the whole population and pie charts showing the proportion of neurons maximally tuned to each parameter value for significantly tuned neurons.

**Extended Data Fig. 8.**
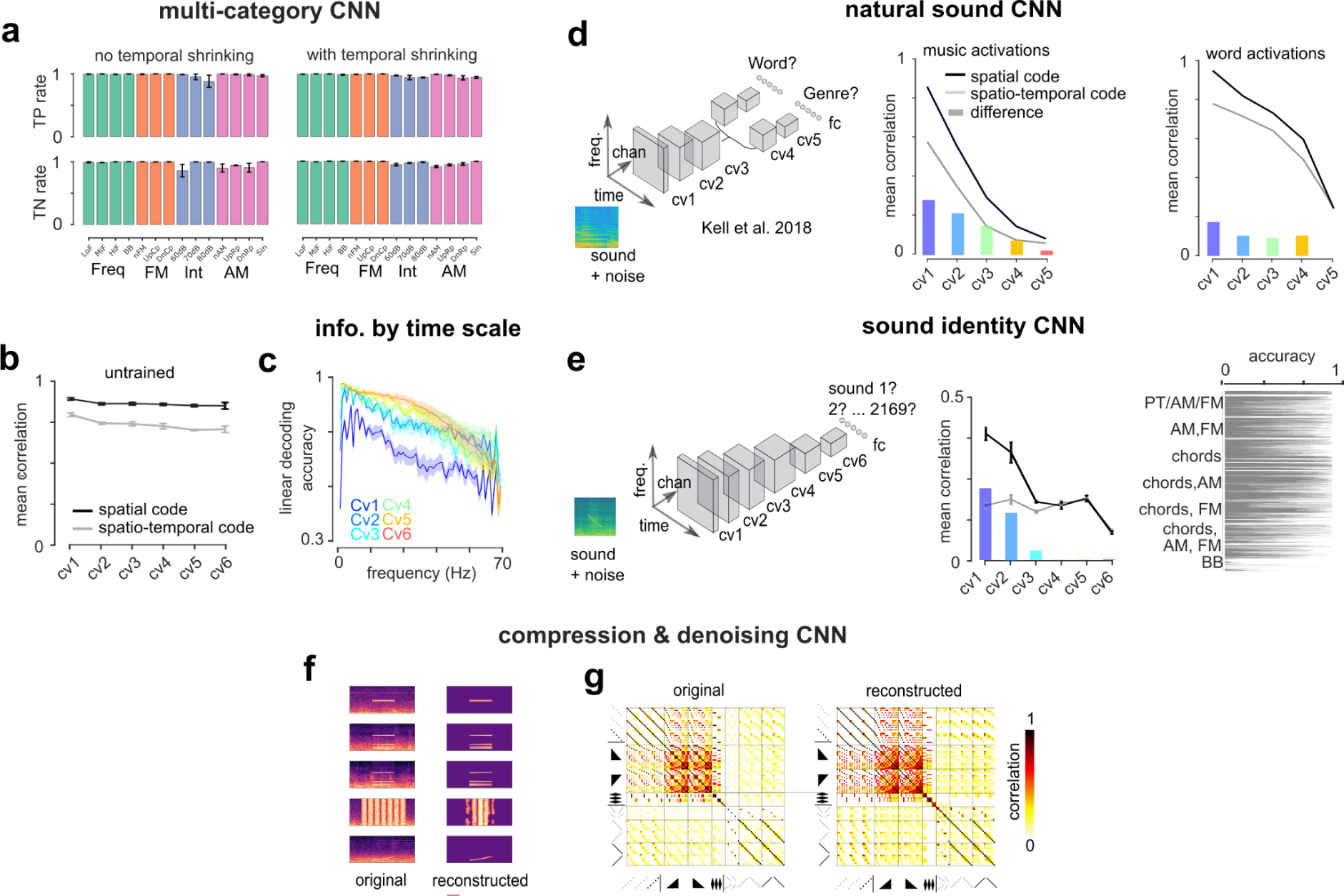
Extended range of CNN networks and details of performance. **a**. Category by category performance of CNNs trained without shrinking of the temporal dimension (left) or with (right) (n=8, error bars are sem). **b.** Mean response correlations from RSA matrices from untrained networks with the same architecture as those trained on the multi-category task (n=8, error bars are sem). **c.** Mean decoding accuracy based on successive Fourier coefficients of CNN responses. 0Hz = spatial code (n=8, shaded areas are sem). **d.** Mean correlations from the network trained on natural sounds from Kell et al ^55^ for musical snippets (left) or words (centre). **e.** Mean correlations from CNNs trained to identify all 2169 sounds individually (left, n=8) and accuracy for each sound (right). **f.** Example sounds provided as input to the autoencoder and their reconstructions at the output. **g.** Representation Similarity Analysis matrix of original sounds and reconstructed sounds showing that the autoencoder fully preserved the relations between all the sounds.

**Extended Data Table 1.** Details of dataset

| *Brain region* | *Recording method* | *Units recorded* | *Responsive units* | *Animal number* | *Session number* | *Units per animal (min, mean, max)* | *Units per session (min, mean, max)* |
| --- | --- | --- | --- | --- | --- | --- | --- |
| Auditory cortex | Cell body 2 photon calcium imaging | 60822 | 19414 (32%) | 7 | 60 | 2164 / 8688 / 20631 | 57 / 1013 / 1782 |
| Auditory thalamus | Axonal bouton 2 photon calcium imaging | 39191 | 3969 (12%) | 4 | 24 | 1280 / 9287 / 19870 | 477 / 1632 / 3120 |
|  | Single unit electrophysiology | 498 | 484 (97%) | 10 | 33 | 4 / 49 / 113 | 2 / 15 / 32 |
| Inferior colliculus | Cell body 2 photon calcium imaging | 15312 | 5936 (39%) | 30 | 101 | 25 / 510 / 2975 | 25 / 151 / 495 |
|  | Single unit electrophysiology | 563 | 442 (78%) | 11 | 30 | 10 / 56 / 119 | 4 / 18 / 54 |

**Extended Data Table 2.**
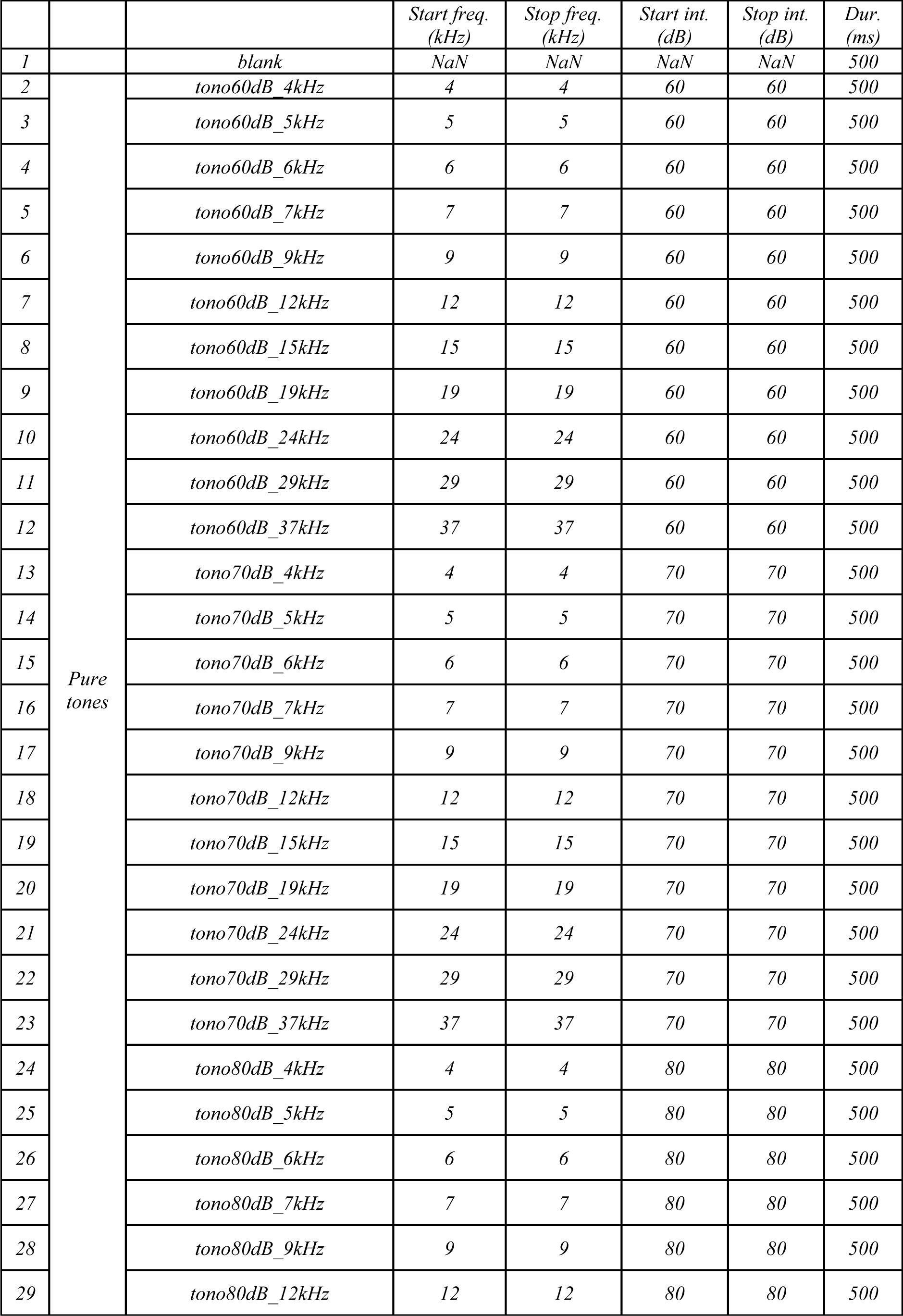
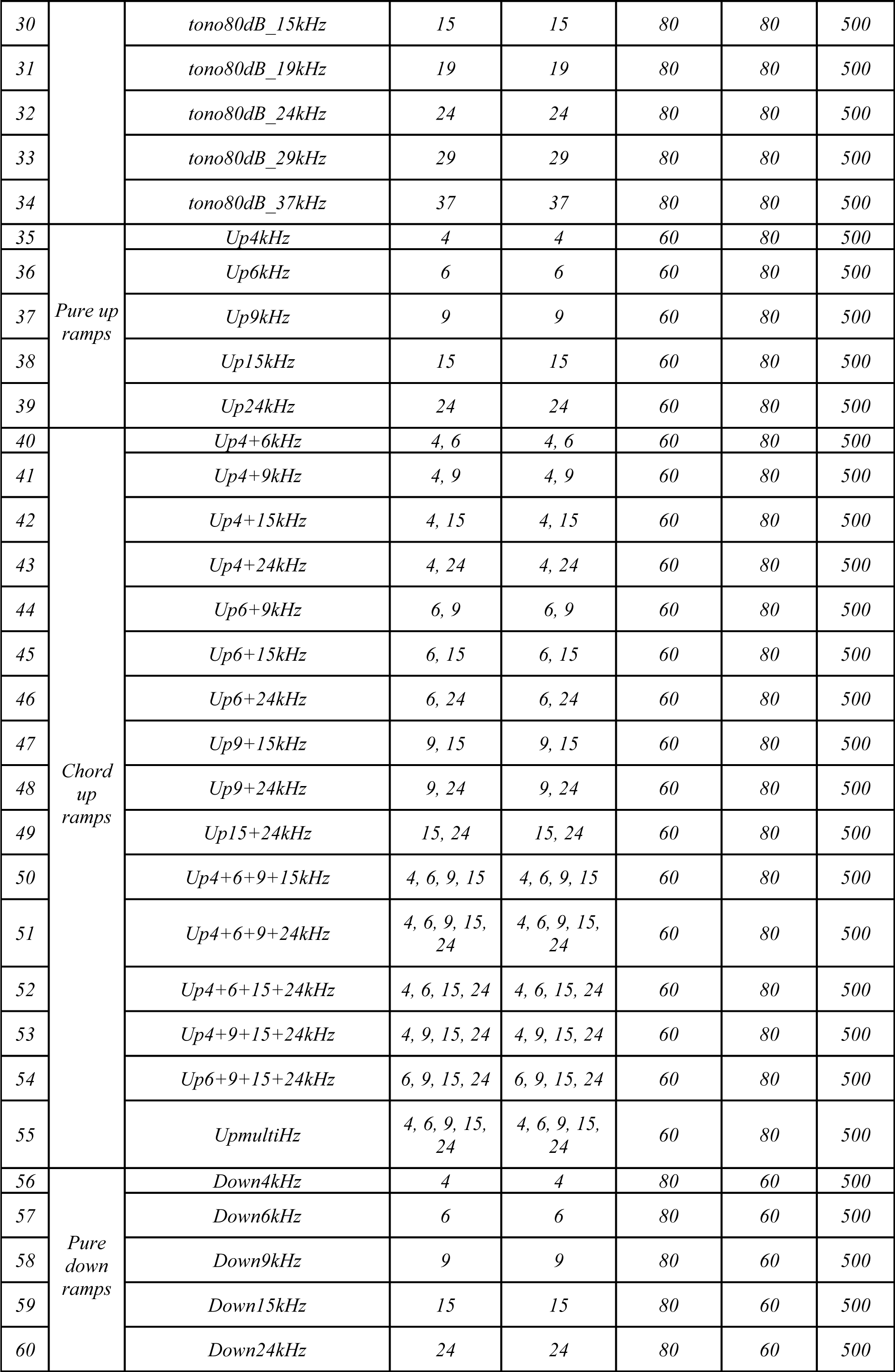
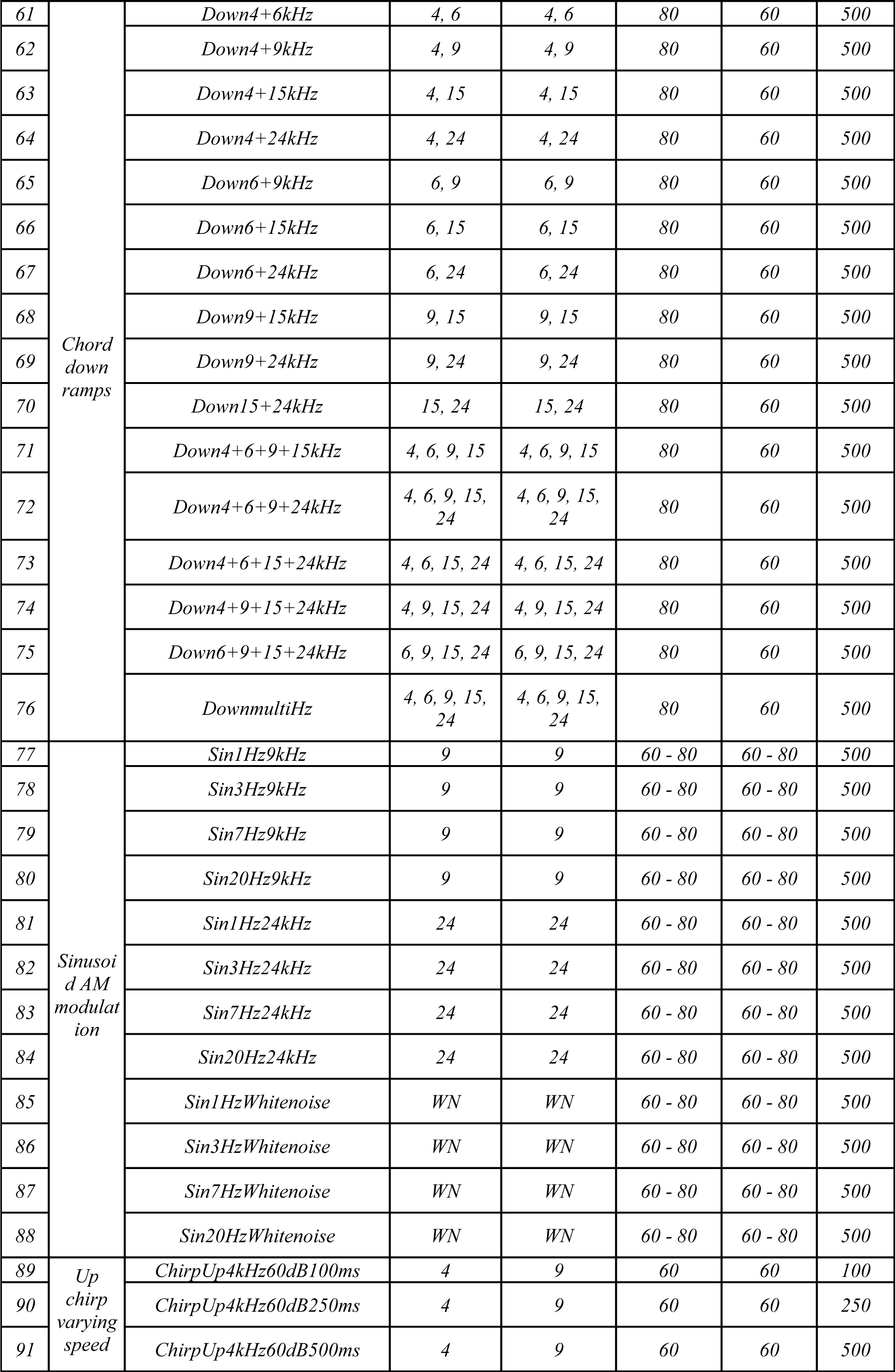
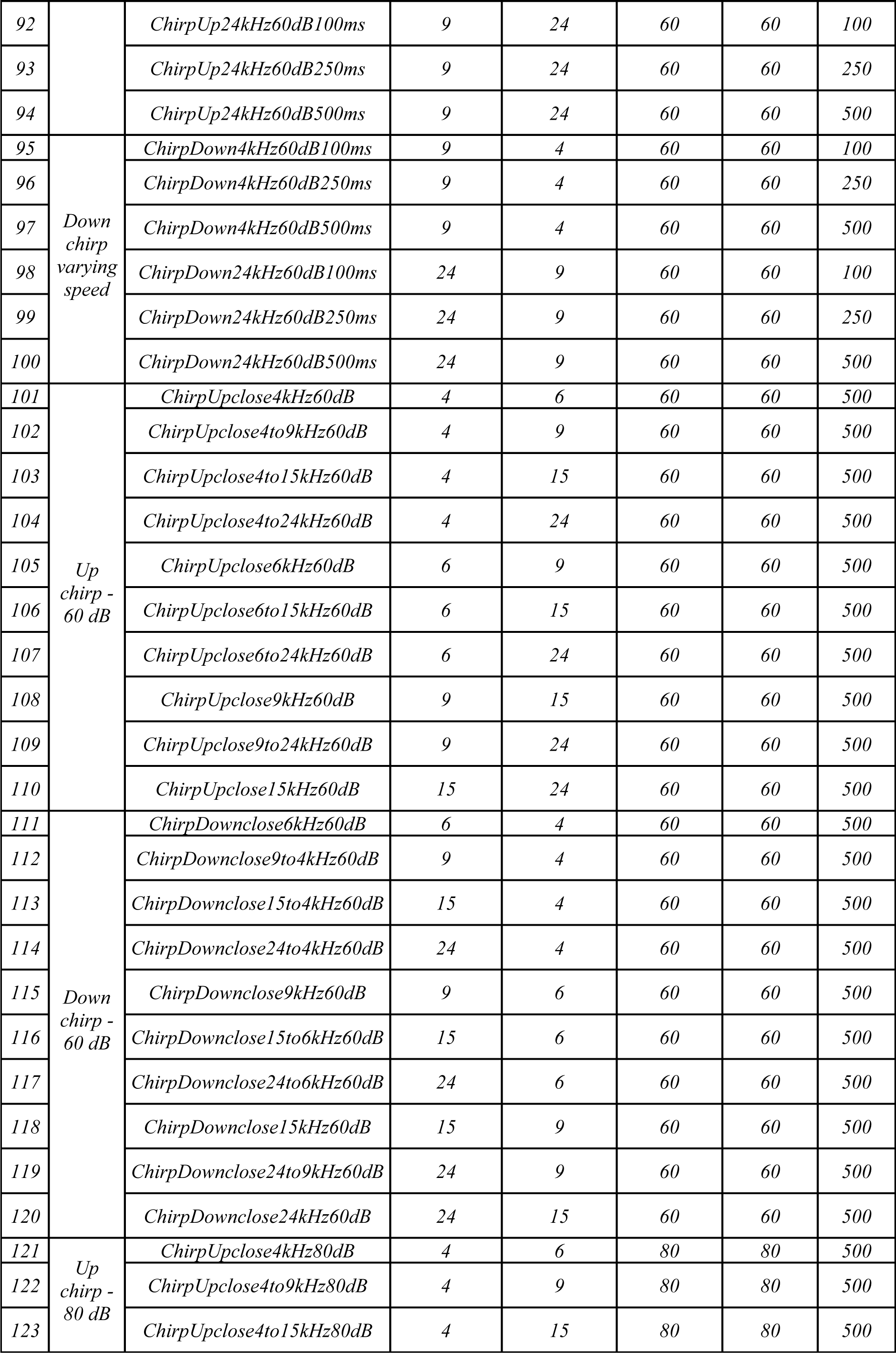
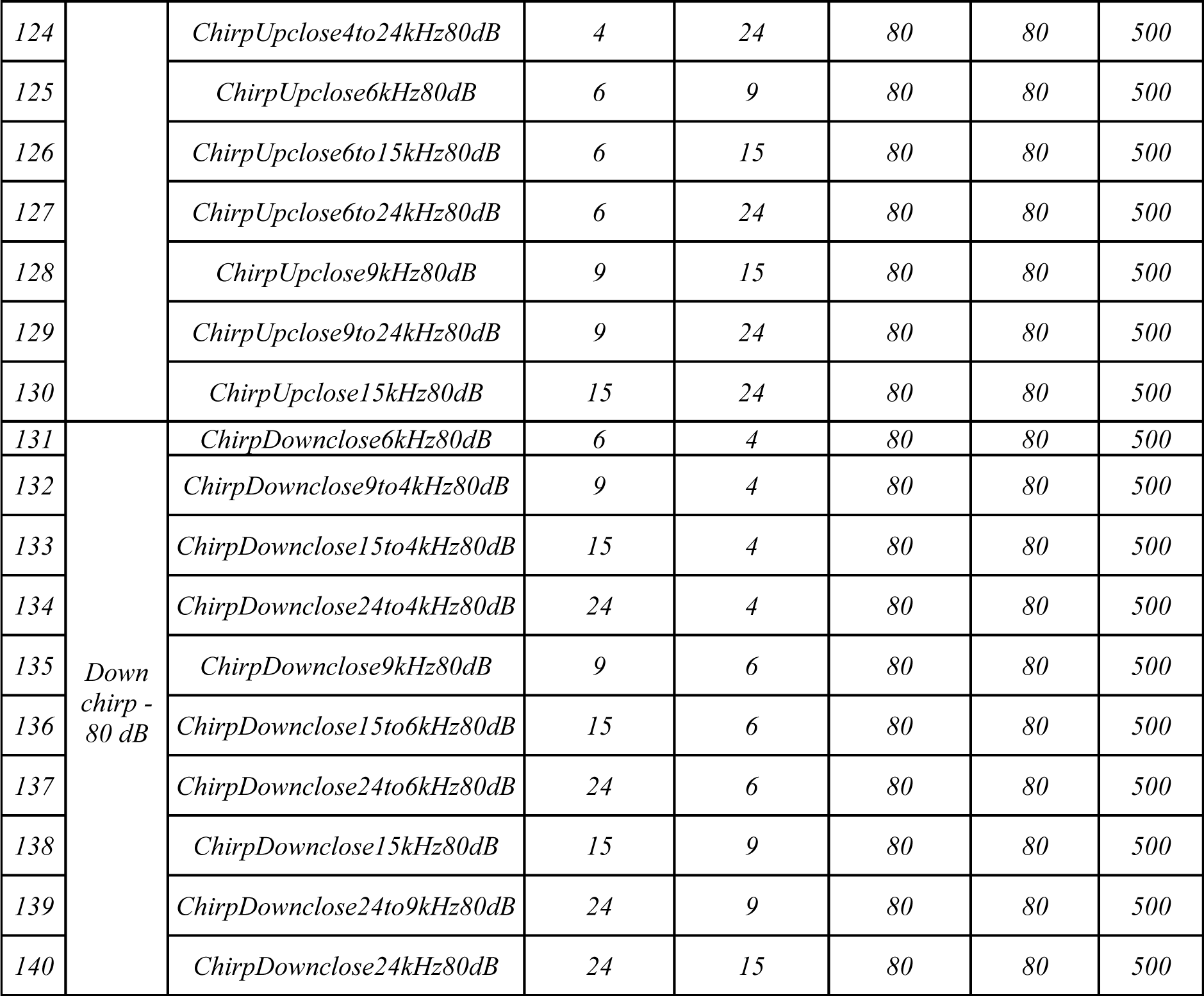
Sound parameters

**Extended Data Table 3.**
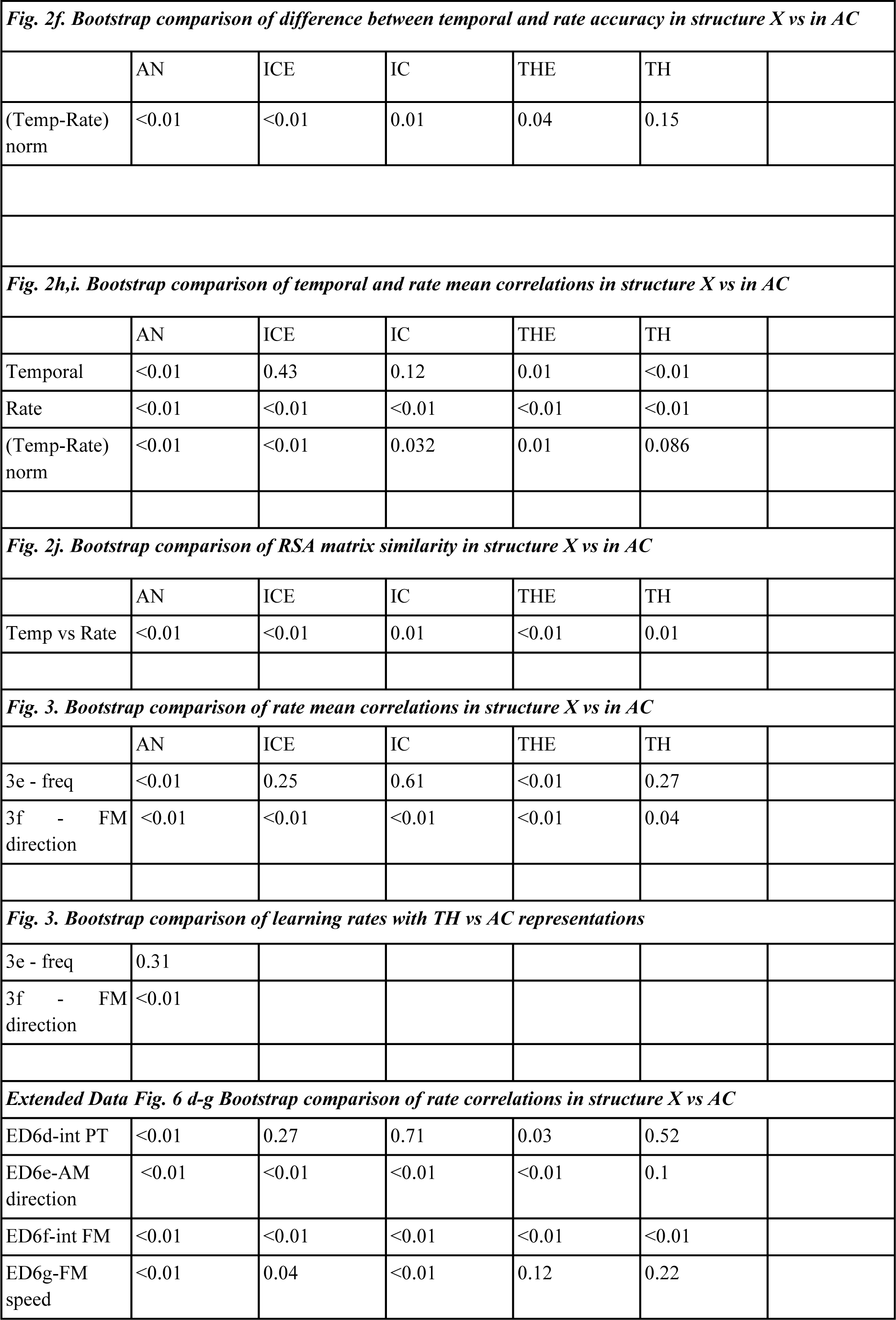
Detail of statistical comparisons

|  |  |  |  |  |  |
| --- | --- | --- | --- | --- | --- |
| **Fig. 2f. Bootstrap comparison of difference between temporal and rate accuracy in structure X vs in AC** |  |  |  |  |  |
|  | AN | ICE | IC | THE | TH |
| (Temp-Rate)<br>norm | <0.01 | <0.01 | 0.01 | 0.04 | 0.15 |
| **Fig. 2h,i. Bootstrap comparison of temporal and rate mean correlations in structure X vs in AC** |  |  |  |  |  |
|  | AN | ICE | IC | THE | TH |
| Temporal | <0.01 | 0.43 | 0.12 | 0.01 | <0.01 |
| Rate | <0.01 | <0.01 | <0.01 | <0.01 | <0.01 |
| (Temp-Rate)<br>norm | <0.01 | <0.01 | 0.032 | 0.01 | 0.086 |
| **Fig. 2j. Bootstrap comparison of RSA matrix similarity in structure X vs in AC** |  |  |  |  |  |
|  | AN | ICE | IC | THE | TH |
| Temp vs Rate | <0.01 | <0.01 | 0.01 | <0.01 | 0.01 |
| **Fig. 3. Bootstrap comparison of rate mean correlations in structure X vs in AC** |  |  |  |  |  |
|  | AN | ICE | IC | THE | TH |
| 3e - freq | <0.01 | 0.25 | 0.61 | <0.01 | 0.27 |
| 3f - FM<br>direction | <0.01 | <0.01 | <0.01 | <0.01 | 0.04 |
| **Fig. 3. Bootstrap comparison of learning rates with TH vs AC representations** |  |  |  |  |  |
| 3e - freq | 0.31 |  |  |  |  |
| 3f - FM<br>direction | <0.01 |  |  |  |  |
| **Extended Data Fig. 6 d-g Bootstrap comparison of rate correlations in structure X vs AC** |  |  |  |  |  |
| ED6d-int PT | <0.01 | 0.27 | 0.71 | 0.03 | 0.52 |
| ED6e-AM<br>direction | <0.01 | <0.01 | <0.01 | <0.01 | 0.1 |
| ED6f-int FM | <0.01 | <0.01 | <0.01 | <0.01 | <0.01 |
| ED6g-FM<br>speed | <0.01 | 0.04 | <0.01 | 0.12 | 0.22 |

