## Supplementary text for "A spatial code for temporal cues is necessary for sensory learning"

### Supplementary Information

Supplementary information about the auditory system dataset

To rapidly obtain large datasets from these structures, we used GCAMP6s-based two-photon calcium imaging of either cell bodies (AC and IC, **Extended Data Fig. 3a & d**) or axonal projections (TH, imaged in AC) (**Extended Data Fig. 3b**). Collecting data simultaneously from around 1000 AC neurons or TH axonal boutons and from 100 to 200 neurons in IC, we could extensively sample representations in each region. In AC, all 60.822 ROIs were mapped to functional subfields based on tonotopic gradients [74] and to the cortical layer from imaging depth (**Extended Data Fig. 4a-f).** 70% of ROIs were in primary auditory cortex (A1), the largest subfield of AC, but the anterior, suprarhinal and dorsal posterior auditory fields were also covered **(Extended Data Fig. 3a & 4e**). Moreover, with recording depth reaching up to 600 µm, we sampled neurons from layers 1 to 5 with an emphasis on layers 2 and 3 (**Extended Data Fig. 4f**). Therefore, with the exception of layer 6 and of the small ventro-posterior subfield, the whole of primary and secondary AC was extensively covered. Inputs from TH were sampled with 39.191 putative TH axonal boutons spread across AC (75% of ROIs in A1) (**Extended Data Fig. 3b**) and validated post-hoc with the thalamic marker VGLUt2 (**Extended Data Fig. 4g,h**)[75]. In addition, we recorded 15.132 ROIs in the dorsal IC down to 250µm depth (**Extended Data Fig. 3d**).

Since calcium imaging and deconvolution has not been verified for TH axons, we performed electrophysiological recording in primary and secondary auditory TH (498 single units, **Extended Data Fig. 3c**). Electrophysiology was also used to cover the central inferior colliculus (563 single units), the main primary subregion of this structure (**Extended Data Fig. 3e**). Electrode locations were identified with post-hoc histology and short-latency responses (**Extended Data Fig. 3c,e**). Finally, we used a detailed biophysical model of the cochlea calibrated against auditory nerve recordings [73] (AN), to provide insight into the information entering the auditory system (**Extended Data Fig. 3f, 4i,j**).

Calcium signals were temporally deconvolved using a linear algorithm to retrieve estimates of neuronal firing rate variations that are robust to parametrization errors [76]. This allowed us to reach a ~150 ms temporal precision as estimated from responses to amplitude modulated sounds (**Extended Data Fig. 3a,b,d**). The temporal modulations of our sounds were chosen to evolve at timescales compatible with this resolution of calcium imaging. This was confirmed by our decomposition of neural population activity into specific timescales using Fourier analysis (**Extended Data Fig. 5e**). This revealed that even with electrophysiology, in which activity contained information at fast timescales up to 30Hz, information nonetheless saturated at around 3Hz. Therefore all information needed to discriminate our sounds is available below 3Hz, which matches calcium imaging resolution, with information at faster timescales being redundant.

Supplementary mathematical derivations

##### ***Population vector notation***

For $N$ neurons indexed $n$, $S$ sounds indexed $s$, $T$ time steps indexed $t$ and $R$ sound repeats indexed $r$, population activity is written:

$$\vec{\nu}_{s,r,t} = (\nu_{1,s,r,t}; ...; \nu_{N,s,r,t})$$

and corresponds to the vector of instantaneous neuronal firing rates for condition $(s, r, t)$. For more compact notations, time averaging is written implicitly:

$$\vec{\nu}_{s,r} = \frac{1}{T}\sum_t \vec{\nu}_{s,r,t}$$

The scalar product of two population vectors is defined as:

$$\vec{\nu}_{s,r,t}.\vec{\nu}_{s,r',t} = \sum_n \nu_{n,s,r,t}\nu_{n,s,r',t}$$

##### ***Estimate of noise-free correlation***

If noise is additive, we can decompose a single-trial population vector for sound $s$ $\vec{\nu}_{s,r}$ as the sum of a noise-free vector $\vec{\nu}_s$ and of an additive normally distributed noise vector $\vec{\xi}_{s,r}$ of mean 0 and variance $\sigma^2/N$. Noise is uncorrelated across trials and sounds. Therefore, in the limit of a large number $N$ of neurons, the scalar product between noise vectors of two different trials $r$ and $r'$ or sounds $s$ and $s'$ converges to zero (i.e. $\vec{\xi}_{s,r}.\vec{\xi}_{s',r'} = \sigma^2\delta_{rr'}\delta_{ss'}$ where $\delta_{rr'}$ is the Kronecker symbol).

For compactness of the demonstration, we assume here that population vectors have zero mean along the neuronal dimension, but the result holds for non-zero mean. The correlation coefficient between two noise-free vectors $\vec{\nu}_s$ and $\vec{\nu}_{s'}$ of mean 0 can be written as:

$$\rho_{\vec{\nu}_s\vec{\nu}_{s'}} = \frac{\vec{\nu}_s.\vec{\nu}_{s'}}{\sqrt{\vec{\nu}_s^{\,2}\vec{\nu}_{s'}^{\,2}}}$$

With trial noise, this becomes:

$$\rho_{\vec{\nu_{s,r}}\vec{\nu_{s',r}}} = \frac{\left(\vec{\nu_s} + \vec{\xi_r}\right)\left(\vec{\nu_{s'}} + \vec{\xi'_r}\right)}{\sqrt{\left(\vec{\nu_s} + \vec{\xi_r}\right)^2\left(\vec{\nu_{s'}} + \vec{\xi'_r}\right)^2}}$$

Because $\vec{\xi_r}$ and $\vec{\xi'_r}$ are two uncorrelated noise vectors, we can write in the limit of large $N$:

$$\rho_{\vec{\nu}_{s,r}\vec{\nu}_{s',r}} = \frac{\vec{\nu}_s.\vec{\nu}_{s'}}{\sqrt{\vec{\nu}_s^{\,2}\vec{\nu}_{s'}^{\,2}}}\frac{1}{\sqrt{(1+\frac{N\sigma^2}{\vec{\nu}_s^{\,2}})(1+\frac{N\sigma^2}{\vec{\nu}_{s'}^{\,2}})}}$$

which can be rewritten as :

$$\rho_{\vec{\nu}_{s,r}\vec{\nu}_{s',r}} = \frac{\rho_{\vec{\nu}_s\vec{\nu}_{s'}}}{\sqrt{(1+\frac{N\sigma^2}{\vec{\nu_s}^2})(1+\frac{N\sigma^2}{\vec{\nu_{s'}}^2})}}$$

If one wants to compare correlation coefficient from different datasets with different unknown $\sigma$'s, this correction factor may introduce discrepancies that are only related to the noise magnitudes.

However, because $\rho_{\vec{\nu}_s\vec{\nu}_s} = 1$ one can derives from the previous equation that:

$$\rho_{\vec{\nu}_{s,r}\vec{\nu}_{s,r'}} = \frac{1}{(1+\frac{N\sigma^2}{\vec{\nu_s}^2})}$$

Combining the last two equations yields in the limit of large $N$:

$$\rho_{\vec{\nu_s}\vec{\nu_{s'}}} \approx \frac{\rho_{\vec{\nu}_{s,r}\vec{\nu}_{s',r'}}}{\sqrt{\rho_{\vec{\nu}_{s,r}\vec{\nu}_{s,r'}}\rho_{\vec{\nu}_{s',r}\vec{\nu}_{s',r'}}}}$$

Note that the formula here is shown only for pairs of trials. In practice, for a finite N, the estimate of the correlation coefficient can be improved by averaging across multiple trial pairs using the formula:

$$\rho_{\vec{\nu}_s\vec{\nu}_{s'}} \approx \frac{\frac{1}{R^2}\sum_{r,r'}\rho_{\vec{\nu}_{s,r}\vec{\nu}_{s',r'}}}{\sqrt{\frac{1}{R^2(1-R)^2}\left(\sum_{r\neq r'}\rho_{\vec{\nu}_{s,r}\vec{\nu}_{s,r'}}\right)\left(\sum_{r\neq r'}\rho_{\vec{\nu}_{s',r}\vec{\nu}_{s',r'}}\right)}}$$

##### ***Noise-corrected sparseness measures***

For simplicity, in this part, we will use $< . >_x$ to indicate averaging across the dimension $x$.

We considered two measures of sparseness, kurtosis and ratio-of- square sparseness *(9, 32, 51, 52)*, which can be applied both for along the neuronal population dimension (population sparseness for a given sound) and the sound response dimension (lifetime sparseness for a given neuron). For $\nu_{n,s}$ the activity of neuron $n$ for sound $s$ averaged across time and trials, the corresponding formula are:

Lifetime sparseness with kurtosis

$$K_n = \frac{< (\nu_{n,s} - < \nu_{n,s} >_s)^4 >_s}{< (\nu_{n,s} - < \nu_{n,s} >_s)^2 >_s^2} - 3$$

Population sparseness with kurtosis

$$K_s = \frac{< (\nu_{n,s} - < \nu_{n,s} >_n)^4 >_n}{< (\nu_{n,s} - < \nu_{n,s} >_n)^2 >_n^2} - 3$$

Lifetime sparseness with ratio of squares

$$S_n = \frac{1 - \frac{<\nu_{n,s}>_s^2}{<\nu_{n,s}^2>_s}}{1 - \frac{1}{S}}$$

Population sparseness with ratio of squares

$$S_s = \frac{1 - \frac{<\nu_{n,s}>_n^2}{<\nu_{n,s}^2>_n}}{1 - \frac{1}{N}}$$

Note that the ratio of square sparseness is properly defined only for positive neuronal activity values.

All these measures, will be biased by variability in the measured responses. To account for this, we express, as above, the response of neuron $n$ to sound $s$ on trial $r$ as:

$$\nu_{n,s,r} = \nu_{n,s} + \xi_{n,s,r}$$

in which $\nu_{n,s}$ is the noise-free response and $\xi_{n,s,r}$ is an additive Gaussian noise term of mean zero and variance $\sigma^2$. In our data, the number of trials is low ($R = 14$) and thus a significant residual noise contribution remains after trial averaging when computing the statistical moments of order larger than 1 that appear in the sparseness measures. For the sake of simplicity, we quantify this here for the moments calculated along the sound dimension (lifetime sparseness). All the reasoning is also valid if moments are calculated along the neuronal dimension (population sparseness) and the estimation will be more robust since number of neurons is much larger than number of sounds.

For the moment of order 1 (average), the noise term vanishes as $1/\sqrt{RS}$ through averaging and is centered on 0 so it will average out across multiple measures:

$$<< \nu_{n,s,r} >_r>_s = < \nu_{n,s} >_s + << \xi_{n,s,r} >_r>_s$$

For the moment of order 2 (variance),

$$<< \nu_{n,s,r} >_r^2>_s = < \nu_{n,s}^2 >_s + 2 < \nu_{n,s} < \xi_{n,s,r} >_r>_s + << \xi_{n,s,r} >_r^2>_s$$

the first noise term is centered on 0 and vanishes as $1/\sqrt{RS}$ but the second noise term represents a positive offset, on average equal to $\sigma^2/R$, which vanishes only as $1/R$, so at least three times less than the first term in our case ($1/\sqrt{RS} = 1/44$ and $1/R = 1/14$). Neglecting all centered terms decaying faster than $1/R$, we thus obtain:

$$<< \nu_{n,s,r} >_r^2>_s = < \nu_{n,s}^2 >_s + \frac{\sigma^2}{R} \tag{1}$$

Noticing that

$$<< \nu_{n,s,r}^2 >_r>_s = < \nu_{n,s}^2 >_s + \sigma^2 \tag{2}$$

we estimate $\sigma^2$ by combining equations 1 and 2:

$$\sigma^2 \approx \frac{R}{R-1}\left(<< \nu_{n,s,r}^2 >_r>_s - << \nu_{n,s,r} >_r^2>_s\right) \tag{3}$$

By combining (1) and (3) we get a noise-corrected estimate of the second order moment of the response by simply averaging over trials before or after taking the squared value:

$$< \nu_{n,s}^2 >_s \approx \frac{1}{R-1}(R << \vec{\nu}_{n,s,r} >_r^2> - << \vec{\nu}_{n,s,r}^2 >_r>_s) \tag{4}$$

Equation 4 can be used to compute the noise-corrected estimates of the ratio of square sparseness using $< \nu_{n,s} >_s^2 \approx << \nu_{n,s,r} >_r>_s^2$.

We now apply this approach to obtain a noise-corrected estimate of ksurtosis. The Kurtosis formula develops into :

$$K_n = \frac{< \nu_{n,s}^4 >_s - 4 < \nu_{n,s} >_s < \nu_{n,s}^3 >_s + 6 < \nu_{n,s} >_s^2 < \nu_{n,s}^2 >_s - < \nu_{n,s} >_s^4}{(< \nu_{n,s}^2 >_s - < \nu_{n,s} >_s^2)^2} - 3 \tag{5}$$

Therefore, in order to correct for noise offsets in the kurtosis, we need the third and fourth order moments of the response. The same approach as used for the second order moments can be used to compute the noise-corrected estimates of the third and fourth moments of the response and of the noise term. The corresponding formula are given here without the details of their derivation:

Third order

$$< \vec{\nu}_{n,s}^3 >_s \approx << \vec{\nu}_{n,s,r}^3 >_r>_s - < \vec{\xi}^3 >_s - 3 < \vec{\nu}_{n,s} >_s < \vec{\xi}^2 >_s$$

$$< \vec{\xi}^3 > \approx \frac{1}{1-\frac{1}{R^2}}(<< \vec{\nu}_{n,s,r}^3 >_r>_s - << \vec{\nu}_{n,s,r} >_r^3>_s + (\frac{3}{R} - 3) < \vec{\nu}_{n,s} >_s < \vec{\xi}^2 >_s)$$

Fourth order

$$< \vec{\nu}_{n,s}^4 >_s \approx << \vec{\nu}_{n,s,r}^4 >_r>_s - < \vec{\xi}^4 >_s - 4 < \vec{\nu}_{n,s} >_s < \vec{\xi}^3 >_s - 6 < \vec{\nu}_{n,s}^2 >_s < \vec{\xi}^2 >_s$$

$$\begin{aligned} < \vec{\xi}^4 >_s \approx & \frac{1}{1-\frac{1}{R^3}}(<< \vec{\nu}_{n,s,r}^4 >_r>_s - << \vec{\nu}_{n,s,r} >_r^4>_s) + \frac{3R-1}{R^3} < \vec{\xi}^2 >_s^2 \\ & + (\frac{4}{R} - 4) < \vec{\xi}^3 >_s < \vec{\nu}_{n,s} >_s + (\frac{6}{R} - 6) < \vec{\xi}^2 >_s < \vec{\nu}_{n,s}^2 >_s \end{aligned} \tag{6}$$

The noise-corrected estimate of kurtosis was therefore computed by replacing the above formula for the moments into equation 5.
